# Root barrier surveillance mechanisms convey soil nitrogen status to shoots

**DOI:** 10.1101/2023.04.19.537470

**Authors:** Defeng Shen, Kathrin Wippel, Simone Remmel, Pengfan Zhang, Noah Kuertoes, Ulla Neumann, Stanislav Kopriva, Tonni Grube Andersen

## Abstract

In roots, formation of the Casparian strip in the endodermal cell walls provides a mechanism for selective uptake of nutrients and water. Establishment of this filter is under spatial surveillance by a receptor-ligand mechanism termed the Schengen pathway. This system provides a mechanism to initiate downstream signaling responses in case of dysfunctional barrier establishment. Due to this interconnected nature, the endogenous physiological role of this survaillance mechanism remain difficult to untangle from the direct barrier function. This is in particularly evident in complex growth setups where multiple inputs are integrated into general whole-plant responses. In this work, we address this by rewiring the genetic network that underlies root barrier formation. We create plants with enhanced, Schengen-independent Casparian strip formation that are not only suitable to probe the role of stronger barrier establishment, but also to evaluate the associated signaling output independently. With focus on the latter, we subjected the rewired plants, as well as a number of established barrier mutants, to multifaceted growth conditions including nitrogen fertilized agricultural soil conditions. By profiling their above- and belowground (a)biotic responses our work reveals that, while increased Casparian strip formation mainly provides the plant with an improved stress resistance, the Schengen pathway is necessary for establishment of a growth-promoting root microbiome and serves to convey information of soil nitrogen status to the shoot. This identifies the Schengen pathway as an essential receptor-based signaling hub for adaptive integration of barrier status, nutritional responses and (a)biotic signaling between above- and belowground tissues.

## Introduction

To survive in highly dynamic environments, plants deploy hydrophobic barriers to protect their vital tissues [1]. One particularly well-studied example is the Casparian strip (CS), which in roots is situated in the endodermal cell layer. The CS blocks water flow within the extracellular matrix and forces solute uptake to occur selectively across the endodermal plasma membrane (PM), analogous to tight junctions in mammals [2]. In most plants, including the model plant Arabidopsis thaliana (hereafter Arabidopsis), the CS consists of circumferentially spanning, lignified “bands” in the anticlinal cell walls, which are formed in the extracellular regions adjacent to the so-called Casparian strip domain (CSD) in the PM [3]. The CSD is established in differentiating endodermal cells by the scaffold-like CASPARIAN STRIP MEMBRANE PROTEINs (CASPs) and a number of secreted enzymes such as the dirigent-like protein ENHANCED SUBERIN 1 (ESB1) responsible for CS fusion in the apoplast [4, 5]. While the mechanisms that ensure a correctly localized CS formation is not completely understood, these enzymes co-localize specifically with the CASP proteins at the CSD and forms a lignification complex in the apoplast. This ultimately ensures deposition of the lignin polymers that constitutes the CS in the middle of the anticlinal cell wall [6].

Expression of most genes involved in CS formation (e.g. CASP1 and ESB1) is controlled by the R3R2 MYB-class transcription factor MYB36 [7, 8]. However, functional barrier establishment further requires an independent signaling mechanism known as the Schengen (SGN) pathway — a surveillance system that co-occurs with the onset of CS establishment [9] and ensures correct fusion of CS components into a coherent barrier [10, 11]. The main components of this system are the stele-synthesized CASPARIAN STRIP INTEGRITY FACTOR (CIF) peptide ligands, their receptor-kinase target SCHENGEN3/GASSHO1 (SGN3/GSO1) and the downstream kinase SCHENGEN1 (SGN1/PBL15) [9, 10, 12]. As CIF peptides diffuse across the unsealed endodermis, they induce activation of SGN3 and SGN1, which results in CS fusion and thereby establishes a self-regulating system by preventing further diffusion of CIF peptides [10]. In mutants where CS function is disrupted (e.g. myb36 knockouts), the resulting increase in CIF diffusion leads to an over-activation of the SGN pathway and initiates a number of mechanisms such as ectopic lignification, induced defense responses and suberin deposition that are able to compensate for the dysfunctional CS establishment [12]. Due to this interconnected nature of the CS and SGN systems, available mutants are limited to those with hyperactive SGN responses or complete absence of both systems [13]. Moreover, as all known mutants have disturbed CS function, the individual physiological role of the CS and the quantitiative, spatially localized, function of the SGN pathway remain difficult to untangle as treatment with CIF peptides can only occur ectopically.

Impaired root barrier formation leads to detrimental growth effects and, under natural conditions, strong changes in the root microbial community composition [13]. However, compensatory responses initiated by the SGN pathway are remarkably effective in compensating abiotic [9, 14] as well as biotic [13] defects, which highlights the importance of this system. Common to the mutants with disturbed function of CS are changes in the ionomic profile — in particular with respect to potassium (K), which is assumed to leak out from the roots due to lack of retention by the CS. However, recent evidence indicates that the SGN system is involved in sensing of the local potassium (K) status via a ROS/Ca2+-dependent signaling mechanism [15]. This suggests a more broad role of the SGN system in systemic integration of nutirent responses. Moreover, mutants with hyperactivated SGN pathway show induced abscisic acid (ABA)-related responses in the shoots [16], which implicates the SGN pathway in a mechanism that convey signals between roots and shoots.

Thus, despite this convincing evidence that the SGN system is physiologically important, most of this work is based on non-functional mutants and we lack deeper insights into how this signaling pathway is integrated at the whole-plant level. In this work, we approach the missing genetic resolution from a semi-synthetic angle by creating and characterizing a new model system with increased, SGN-independent CS formation. Through an experimental setup with focus on different soil conditions, we asked the question if the SGN pathway is an important signaling hub for integration of nutritional responses between above- and belowground tissues. Moreover, as we employed this under natural soil conditions where plants maintain a functional microbiota, we were further able to put this into a complex multifaceted (a)biotic stress situation that represents agriculturally relevant situations.

## Results

### Rewiring MYB36 creates an earlier and SGN-independent Casparian strip

To create plants with SGN-independent CS, we employed the promoter region of the direct MYB36 target CASP1 [8] to drive MYB36 expression. In principle, this generates a spatially restricted feedback overexpression where MYB36 induces itself and its downstream targets in cells that are already initiating CS formation (MYB36Loop). To evaluate functionality of this, we performed a whole-root transcriptome analysis of homozygous lines carrying the MYB36Loop construct and compared them to the *myb36-2* mutant and the respective parental lines. Indeed, presence of MYB36Loop led to strongly increased expression of both MYB36 and CASP1, which were almost non-detectable in the *myb36-2* mutant (Fig. 1A). Within our significance threshold (false recovery rate (FDR) < 0.05, |Log2 FC |> 2), a subset of 113 differentially expressed genes (DEGs) showed a similar response as *CASP1* and *MYB36*, whereas 142 showed the opposite behavior (i.e repressed in MYB36Loop and induced in *myb36-2*) (Fig. S1A). Within the first set, we found most genes with a characterized function in CS establishment (including *SGN1* and *SGN3*) (Fig. 1B), which indicates that the MYB36Loop activates the CS-forming machinery. In line with this, the functional gene ontology (GO) term “cell-cell junction assembly” was overrepresented among the genes induced in MYB36Loop and repressed in *myb36-2*, respectively (Fig. S1B). Plants expressing MYB36Loop had an increase in CS width from ca. 500 nm in WT to an average of approximately 1500 nm (Fig. 1C-D), which confirms that this transcriptional response led to a post-translational output. Moreover, MYB36Loop lines also exhibit earlier onset of a functional CS formation (Fig 1E). One additional observation was that the SGN-induced genes *PER15/49* and *MYB15* [11, 14] were not affected in MYB36Loop (Fig. 1B) and GO terms associated with an activated SGN system were repressed in MYB36Loop (Fig. S1A-B). One CS-related gene, *CASP4* was even repressed in the MYB36Loop lines (Fig. 1B) and appear to require both MYB36 and SGN for expression (Fig. S1C). This indicates that in MYB36Loop plants, the early onset CS may override SGN activation directly by preventing diffusion of CIF peptides. To substantiate this, we introduced the MYB36Loop construct into *sgn3-3* mutants. As this could fully complement the defective CS formation found in this line [9] (Fig. 1F-G), we concluded that the ectopic expression of MYB36 in the MYB36Loop lines uncouples CS barrier formation from its endogenous SGN-dependent program. MYB36Loop plants did not show any significant change in xylem lignification onset nor ectopic CS formation in adjacent cell types (Fig. S1D-E).

**Figure 1.**
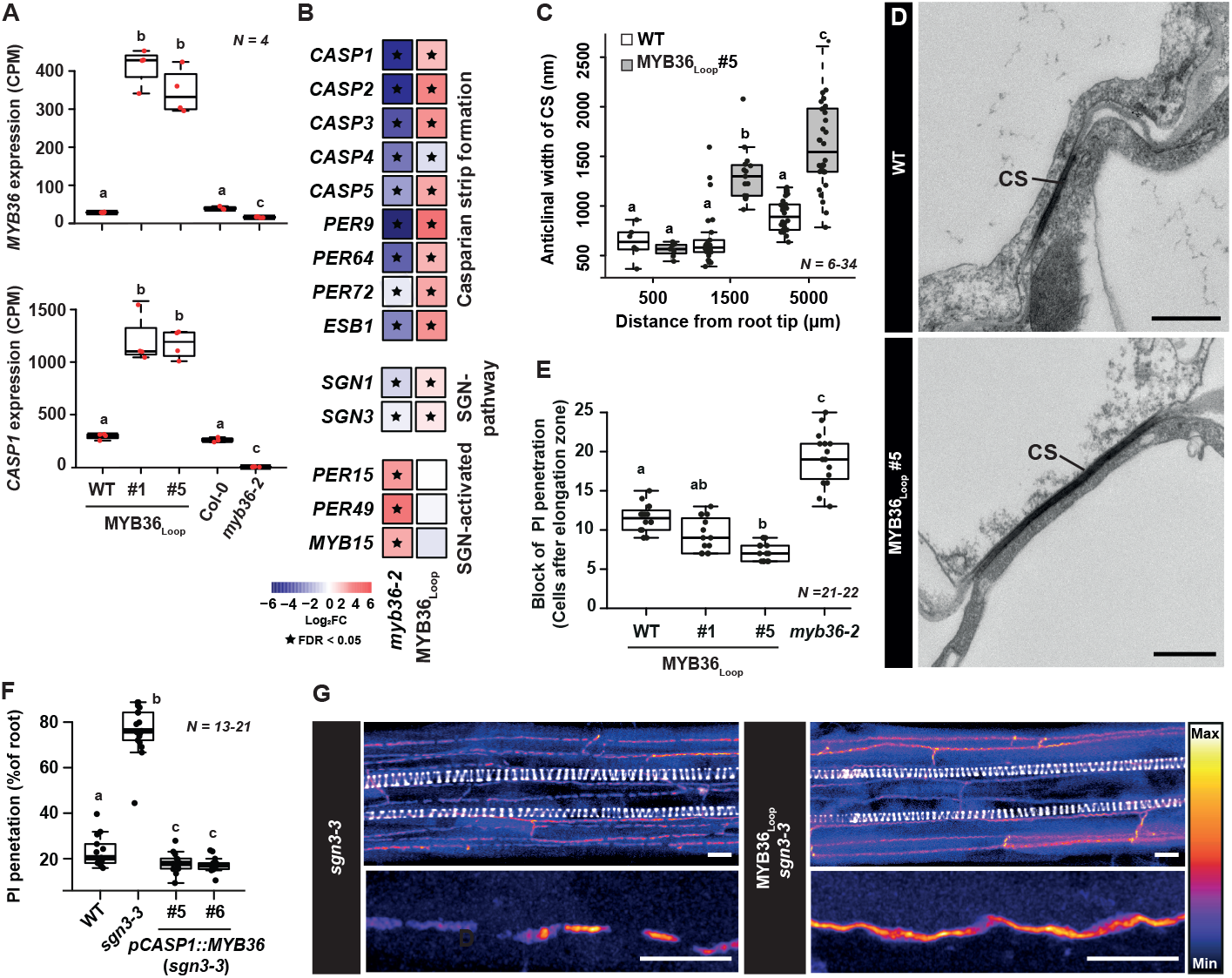
Transcriptional responses and Casparian strip formation in plants expressing MYB36 driven by the CASP1 promoter. (A) Normalized read counts of MYB36 and CASP1 (counts per million, CPM). (B) Heatmap depicting expression of individual genes from transcriptome analysis of whole roots. Star symbols represent significantly differentially expressed (FDR < 0.05) genes compared with the corresponding parental lines. (C) Quantification of Casparian strip width in transmission electron microscopy (TEM) images. (D) TEM images of Casparian strips (pointed by black lines) between two endodermal cells, scale bar: 500 nm. (E) Quantification of onset of propidium iodide blockage. (F) Measurement of the percentage of roots that can be penetrated by propidium iodide (PI). (G) Maximum projection of a confocal image stack of Basic Fuchsin-stained 7-day-old roots. Scale bars in upper images represent 10 □m whereas scale bars in lower images represent 5 □m. For boxplots, the center line indicates median, dots represent data points, the box limits represent the upper and lower quartiles, and whiskers maximum and minimum values. Letters depict statistical difference in a one-way ANOVA analysis with a Holm-Sidak-adjusted posthoc t-test (P < 0.05); CS, Casparian strip.

### Overloading the Casparian strip domain leads to ectopic CS-like structures

To investigate the dynamics of CS establishment, we generated the MYB36Loop lines in a pCASP1::CASP1-GFP background [3]. Onset of CASP1-GFP expression follows a “string-of-pearls” pattern in the anticlinal PM of differentiating endodermal cells [3] (Fig. 2A). However, in MYB36Loop plants, approximately 2 hours after onset, the CASP1-GFP signal quickly split into two laterally expanding lines that likely represent the increased CS width (Fig. 1C-D). Moreover, after about 7 hours the CASP1-GFP signal formed “ring-like” radially expanding patches in the periclinal cell walls (Fig. 2A-B and Movie S1). These structures contained lignin-specific signals (Fig. 2B), which indicates that the entire lignification program responsible for CS polymerization was also mis localized to these patches. To test this, we created combinatorial lines that expressed the MYB36Loop construct, fluorescent markers for CASP1-GFP and the lignifying enzymes ESB1-mCherry or PER64-mCherry. Indeed, these enzymes co-occurred with the ectopic CASP1-GFP patches, although in a broader zone that extended beyond the CASP1-GFP signal (Fig. 2C and Fig. S1F). In combination with the expanding nature of the CASP1-GFP signal (Movie S1), this suggests that lignin polymerization may occur at the edge of these patches and be necessary for their expansion in the periclinal wall. In line with this, plants treated with the monolignol synthesis inhibitor piperonylic acid (PA) [17, 18] failed to form coherent patches (Fig. 2D). Moreover, when introducing the MYB36loop construct into the *esb1-1* background with a fluorescent marker for CASP1-GFP [5], we found no ectopic patch formation, but rather vesicle-like structures that appeared to be located in the cell lumen (Fig 2E). Interestingly, MYB36Loop plants had a slight delay of endodermal suberization (Fig. S1H) and that treatment with 100 nM CIF2 had no effect on the suberin patterning in MYB36Loop lines (Fig. S1H-I). Moreover, suberin was excluded from the lignified patches within the individual cells (Fig. S1G). Combined, our findings support a model where feedback-driven MYB36 expression overloads the CSD with components of the CS-forming machinery. As a consequence, each endodermal cell creates a wider CS and has spill-over deposition of CS-machinery elements into the periclinal PM and cell walls (Fig. S1J). Through an ESB1-dependent mechanism, these enzymes form “islands” of CS-like structures that mimic the CS formation and inhibits suberization.

**Figure 2.**
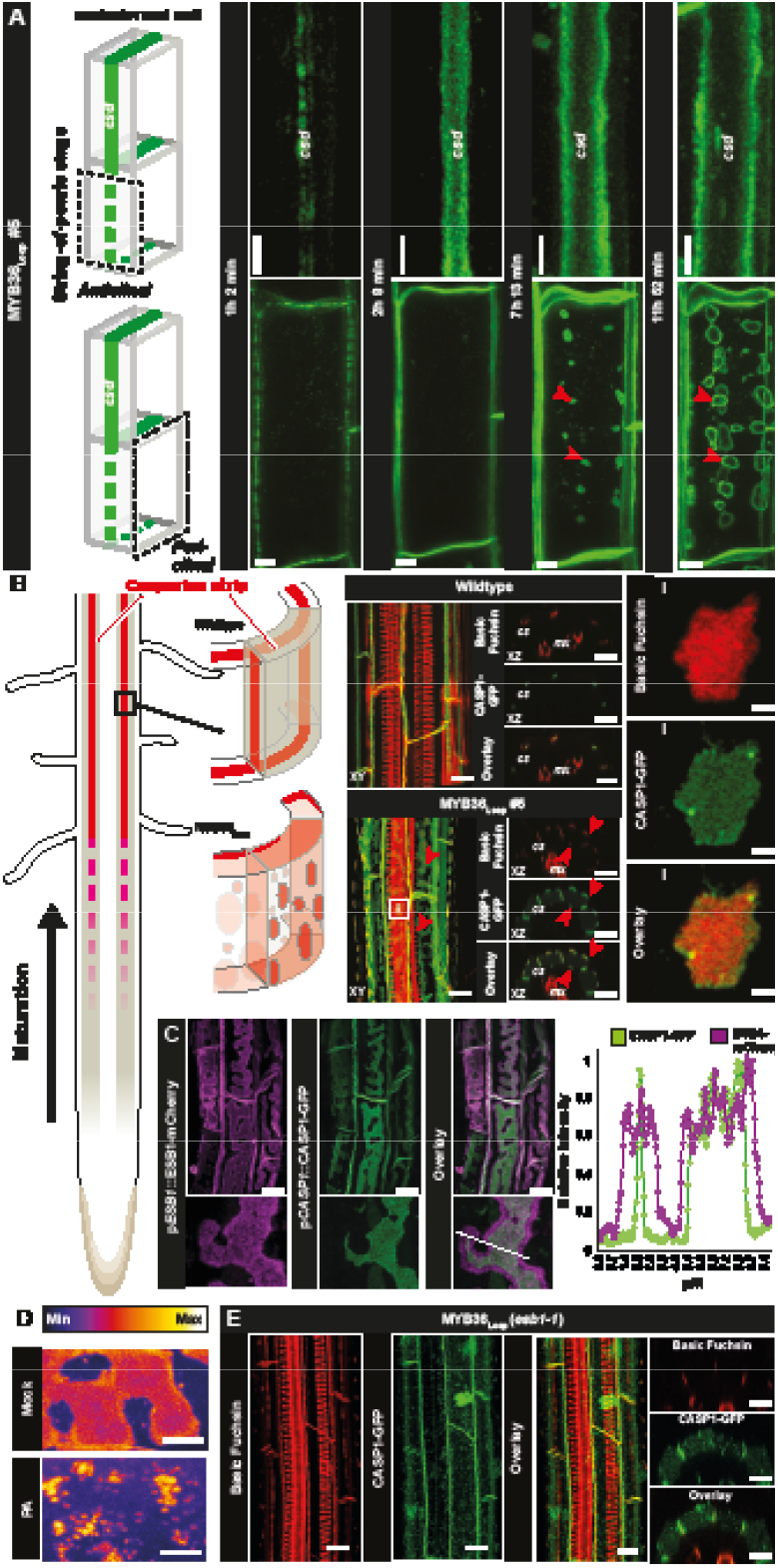
Analysis of Casparian strip onset and Schengen responses in plants expressing MYB36 driven by the CASP1 promoter. (A) Time course of CASP1-GFP expression in MYB36Loop#5 plants. Imaging was initiated at the string-of-pearls stage before onset of CASP1-GFP expression. Images are a maximum projection of confocal image stacks from the anticlinal (upper graph) and periclinal (lower graph) view of an individual endodermal cell. Casparian strip domain (CSD), red arrowheads point to ectopic CASP1-GFP in the periclinal plasma membrane. (B) Maximum projections of confocal image stacks of Basic fuchsin-stained wildtype and MYB36Loop#5 roots expressing the CASP1-GFP fusion reporter in the region where mature CS is formed. Arrowheads represent ectopic lignin-specific signals on the periclinal cell walls of endodermis in MYB36Loop#5 roots. For the insert (i) scale bar: 2 µm. (C) Maximum projection of a confocal image stack of Basic fuchsin stained 7-day-old MYB36Loop roots expressing pESB1::ESB1-mCherry and pCASP1::CASP1-GFP. Line in overlay depicts the transect used for relative intensity measurements. (D) Periclinal ectopic CASP1-GFP depositions in MYB36Loop#5 roots treated with mock or 10 □M piperonylic acid (PA) for 48 hours. Scale bars: 2 µm. (E) Maximum projection (left) and cross-sectional view (right) of a confocal image stack of Basic fuchsin-stained 7-day-old esb1 roots expressing pCASP1::CASP1-GFP and MYB36Loop. Unless otherwise stated, the scale bars represent 10 µm.

### Rewired Casparian strip formation provides increased abiotic stress resistance

*Next, we set out to evaluate how the MYB36Loop r*oots respond to abiotic stress factors. Under standard agar-based conditions MYB36Loop lines showed a slight but significant reduction in primary root length (Fig. 3A, Fig. S2A). This was accompanied by a strong reduction in lateral root (LR) density — similar to what has been described in *myb36-2* (Fig. 3B, Fig. S2A) [19]. Yet, MYB36Loop lines had an earlier repression of primordia development than *myb36-2* (Fig. S2B) and these appeared “flattened” against the endodermis (Fig. S2C), most likely due to the increased CS deposition creating a mechanical hindrance for root emergence. When subjected to salt or osmotic stress, MYB36Loop plants had improved primary root growth compared to their parental line (Fig. 3C), which is consistent with previous observations which demonstrated that CS deposition can occur earlier in order to cope with increased stress [13, 15]. Remarkably, under nutrient-rich (½ MS) conditions, MYB36Loop roots displayed increased expression of genes encoding for nitrogen/phosphorus-related stress responses such as NIN-LIKE-Proteins (NLPs) [20] as well as a reduction in repressors belonging to the NITRATE-INDUCIBLE, GARP-TYPE TRANSCRIPTIONAL REPRESSORs (NIGTs) [21] and BTB AND TAZ DOMAIN PROTEINs (BTs) families [22] (Fig. 3D). This suggests that misalignment of the CS/SGN systems affects signaling associated with nitrogen homeostasis in the root. When grown in absence of nitrogen (½ MS agar medium without added nitrogen), no significant changes were observed in *myb36-2* plants, whereas MYB36Loop plants showed increased relative root growth when compared to its parental line (Fig. 3E). No such changes were observed in the size of the shoots, however a similar root response occurred when phosphorus was omitted, whereas this was not the case under sulfur-depleted conditions (Fig. S2D). This suggests that the improved root growth in MYB36Loop plants is likely due to changes in nitrogen/phosphorus signaling in accordance with the transcriptional responses. While the specific role of nitrogen in root barrier establishment remains unclear, CS has recently been found to respond to nitrogen status in maize (Zea mays) [23]. Therefore, we analyzed if the establishment of a functional CS responds to nitrogen-restrictive conditions in Arabidopsis. Remarkably, WT plants showed a dose-dependent delay of CS function when nitrogen supply was restricted. However, this was not observed in *myb36-2* nor in MYB36Loop plants (Fig. 3F). Taken together, we propose that earlier apoplastic blockage confers increased resistance to abiotic stresses and highlights the importance of correct spatial alignment of the CS-SGN in particular for nitrogen-related signaling.

**Figure 3.**
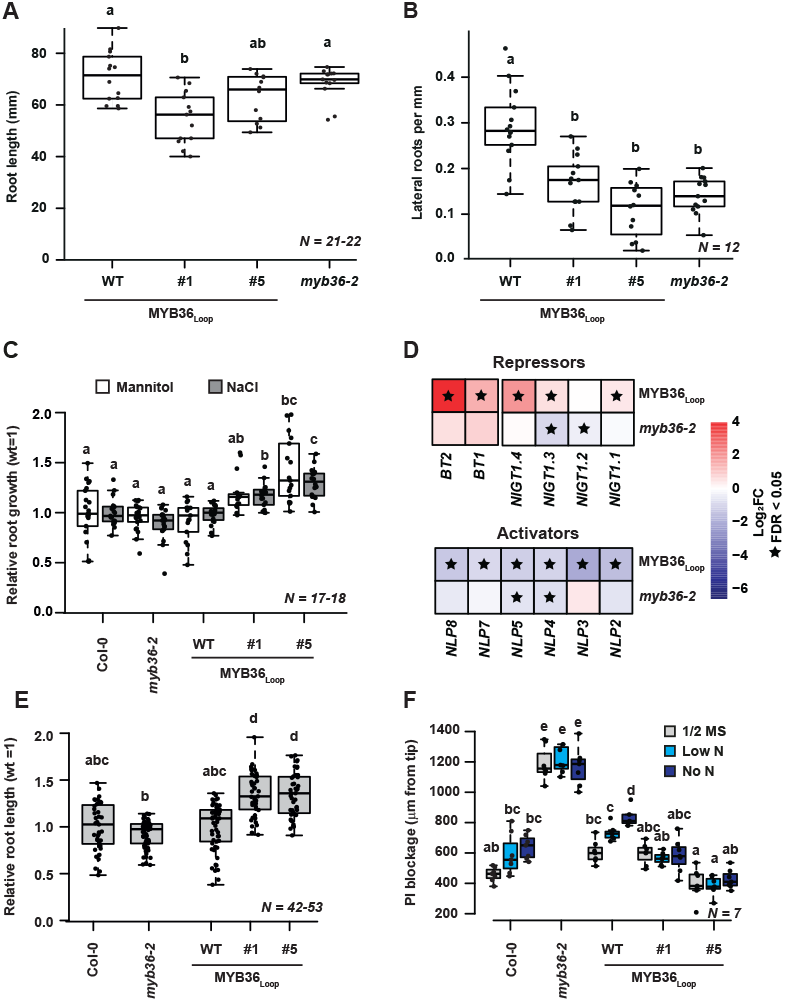
Root architecture and abiotic stress analysis in mutants with modified Casparian strip formation. (A) Primary root length of 14-day-old plants from ½ MS agar. (B) Lateral root density of 14-day-old plants grown on ½ MS agar. (C) Relative root growth of 14-day-old plants after 7 days under stress conditions (150 mM mannitol or 100 mM NaCl). (D) Heatmap of genes belonging to the BTB AND TAZ DOMAIN PROTEINs (BT), NITRATE-INDUCIBLE, GARP-TYPE TRANSCRIPTIONAL REPRESSORs (NIGT) or NIN-LIKE Protein (NLP) families in roots of MYB36Loop and myb36-2 plants. Star symbols represent significantly differential expressed (FDR < 0.05) genes. (E) Relative root growth of 14-day-old plants grown without added nitrogen for 11 days. (F) Onset of functional Casparian strip by means of propidium iodide (PI) blockage. ½ MS agar medium (½MS), 0.11 mM N (Low N) or 0 mM N (No N). For boxplots, the center line in the box indicates median, dots represent data, limits represent upper and lower quartiles, and whiskers maximum and minimum values. Different letters depict statistical difference in a one-way ANOVA analysis with a Holm-Sidak-adjusted posthoc ttest (P < 0.05). #1 and #5 refer to two independent homozygous lines expressing the pCASP1::MYB36 construct in the pCASP1::CASP1-GFP background (WT).

### Misalignment of Schengen activation impairs establishment of beneficial plant-microbe interactions in roots

With a clear influence of the MYB36Loop on abiotic root responses established, we set out to create a more complex environment for the root and assess if establishment of beneficial microbe associations is affected in MYB36Loop plants. In contrast to previous studies that have investigated the impact of barrier formation on naturally occurring bacterial communities [13], we initially used a gnotobiotic system with a synthetic bacterial community (SynCom). The purpose of this particular setup was to readout functional plant-microbe interaction in the form of induced plant growth promotion (PGP) [24]. After five weeks of growth, both parental lines and the *myb36-2* mutant showed a significant increase in rosette fresh weight in presence of the SynCom (Fig. 4A). This is consistent with previous observations indicating that the *myb36-2* mutant retains the ability to establish a healthy microbiome despite having a defective barrier [7, 8]. Surprisingly, PGP was completely abrogated in MYB36Loop plants (Fig. 4A). This indicates that either SGN-related signaling is essential in establishing beneficial microbe associations or the increased barrier function in the MYB36Loop lines impairs this. Across all used lines, we found only minor changes in the relative abundance of the employed bacterial strains (Fig. 4C, Fig. S3A-D), which indicates that the lack of PGP in MYB36loop was not due to disturbances in the ability to shape the bacterial community. However, the bacterial load was strongly increased in the *myb36-2* plants, but decreased in MYB36Loop (Fig. 4B). Thus, in this simplified setup, root colonization rather than community composition is important for PGP and that this is controlled by correct coordination of the CS-SGN system. To investigate how this translates into naturally occurring, uncultured microbial communities, we grew MYB36Loop#5 (which overall displayed a stronger effect than #1) and the *myb36-2* mutant in our local soil [25] (Cologne Agricultural Soil, CAS), and profiled both bacterial and fungal communities associated with the root and the rhizosphere. In contrast to the gnotobiotic setup, MYB36Loop roots had an increased bacterial and fungal load, which, surprisingly, did not occur in *myb36-2* (Fig. 4D). The alpha diversity of bacterial and fungal communities was strongly reduced in both roots and the rhizosphere of MYB36Loop (Fig. 4E). Combined with the increased microbial load in the root, this may indicate that opportunistic species have an advantage to colonize the vicinity of MYB36Loop plants. Analysis of the beta-diversity revealed divergent microbial communities in soil, rhizosphere, and root samples (Fig. S4A and C) and significantly different community structures on roots according to genotype (Fig. S4B and D). The strong decrease in the diversity of microbial communities were characterized by prominent changes in the relative abundance of several bacterial and fungal families across the measured compartments, particularly in the MYB36Loop expressing plants (Fig. 4F, Fig. S4E and G). This further points to an enhanced proliferation of opportunistic species in these plants under natural conditions. Combined, we propose that spatial coordination of the CS and the associated SGN signaling has strong influence on establishment and control of beneficial biotic interactions in the root. Under natural conditions, spatial misalignment of these systems creates a suboptimal niche environment in the rhizosphere that allows increased growth and colonization of opportunistic microbes.

**Figure 4.**
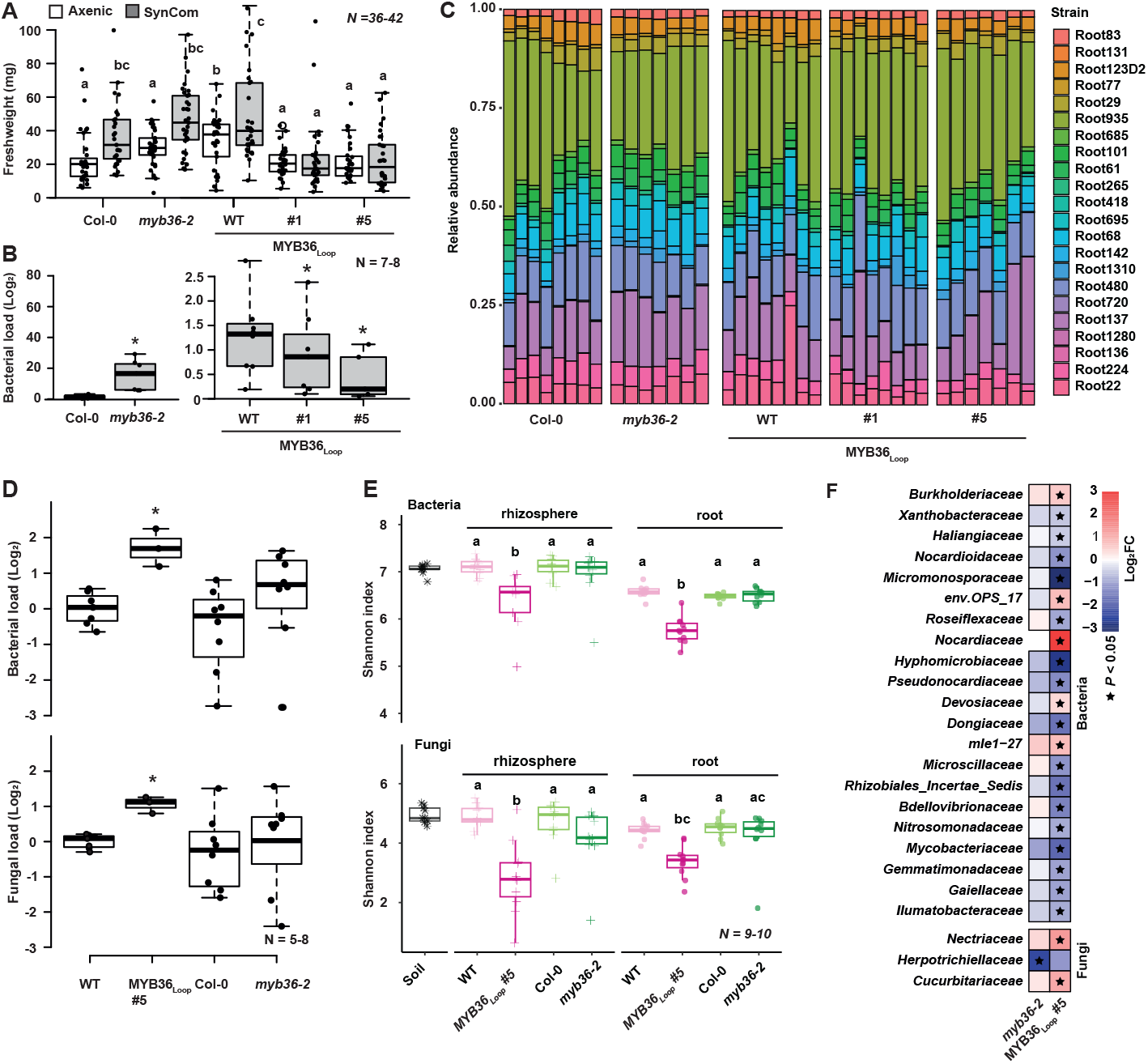
Analysis of synthetic and naturally occurring microbial communities in roots. (A) Fresh weight of 5-week-old rosettes with mock treatment (Axenic) or a synthetic bacterial community (SynCom). Letters depict statistical difference in a one-way ANOVA analysis with a Holm-Sidak-adjusted posthoc ttest (P < 0.05). (B) Bacterial load of plants co-cultivated with the SynCom. Asterisks depict significant changes according to a two-sided Student’s t-test vs parental line (P < 0.05). (C) Relative abundance of bacteria in roots of 5-week-old plants. (D) Bacterial (upper graph) or fungal (lower graph) load of 4-week-old plants grown under CAS conditions. (E) Alpha diversity of bacterial (upper graph) and fungal (lower graph) communities in soil, root and rhizosphere of pCASP1::CASP1-GFP (WT), MYB36Loop#5, Col-0, and myb36-2 plants grown in CAS. Letters indicate significant differences according to pairwise Wilcoxon rank test with Bonferroni correction. (F) Heatmap depicting bacterial and fungal families in mutant roots compared to corresponding parental lines, based on Log2-transformed fold changes. Stars indicate significant (Wilcoxon test with Bonferroni correction, adjusted P < 0.05). For boxplots, center line indicates the median, dots represent data, box limits upper and lower quartiles, whiskers maximum and minimum values. #1 and #5 refer to two independent homozygous lines expressing the pCASP1::MYB36 construct in the pCASP1::CASP1-GFP background (WT).

### Increased Casparian strip deposition disturbs shoot mineral content in a soil dependent manner

To test if different soil types would establish different CS-associated responses in the shoots, we set out to investigate the effects of MYB36Loop shoot performance under different soil conditions. For this, we used standard potting soil to represent a nutrient-rich environment and compared this to CAS, which naturally has a low nitrogen content [25]. Independent of the soil type, *myb36-2* and MYB36Loop rosettes were significantly smaller than their parental lines (one-way ANOVA, P < 0.05) (Fig. 5A, Fig. S2E). As barrier mutants show characteristic changes in their shoot ionome [13], we analyzed the mineral ion content of shoots across the two employed soil types. MYB36Loop and *myb36-2* rosettes accumulated distinct mineral profiles when compared to their parental lines in both soil conditions (FDR < 0.05) (Fig. 5B, Fig. S5A). However, about 78% of variation in mineral content could be explained by the soil type exceeding the proportion explained by the genotype (Fig. 5B). The strongest effects were observed when plants were grown on CAS, where most heavy metals were increased in MYB36Loop plants but decreased in *myb36-2* (Fig. 5C, Fig. S5A). This proposes that the opposing CS status in these lines underlie the contrasting mineral profiles. Yet, independent of soil type, both MYB36Loop and *myb36-2* rosettes had significantly decreased amounts of potassium (K) (one-way ANOVA, P < 0.01) (Fig. 5C, Fig. S5A). Low K accumulation in shoots consistently correlates with ineffective barrier formation [9, 13] and it was therefore unexpected to find a reduction in K content in MYB36Loop plants. However, this reduced K content might be caused by a deficiency of a SGN-associated K-status signaling mechanism rather than by a direct barrier effect [15]. Correspondingly, the repressed SGN activation in the MYB36Loop plants may prevent proper K-status signaling, whereas in *myb36-2* this reduction is likely due to leakage through a dysfunctional CS.

**Figure 5.**
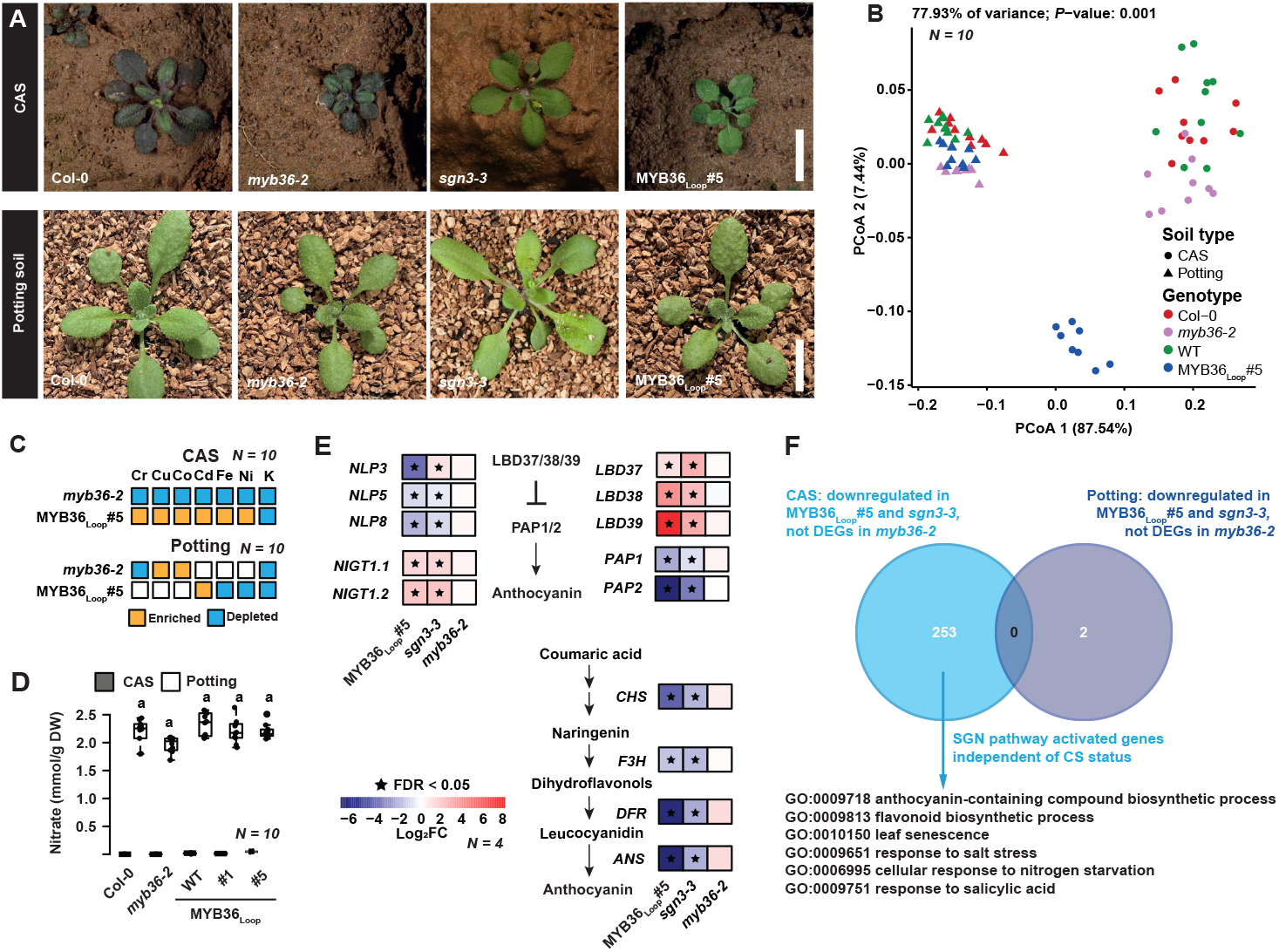
Shoot analysis of plants with modified root barriers grown in different soil conditions. (A) Plants grown on Cologne agricultural soil (CAS) or potting conditions (Potting). Scale bars: 1 cm. (B) PCoA plot of Bray-Curtis distances calculated on 24 shoot mineral contents of 4-week-old plants grown under CAS or standard potting soil conditions. 77.93% variation in shoot ionome can be explained by soil type (P=0.001, PERMANOVA). (C) Minerals and (D) nitrate in rosettes grown on CAS or potting soil conditions. Minerals are normalized to corresponding parental line (two-sided Student’s t-test, P < 0.05). (E) Heatmap showing transcriptional behavior of NIN-LIKE-Proteins (NLPs), NITRATE-INDUCIBLE, GARP-TYPE TRANSCRIPTIONAL REPRESSORs (NIGTs), BTB AND TAZ DOMAIN PROTEINs (BTs) families, PRODUCTION OF ANTHOCYANIN PIGMENT 1 and 2 (PAP1 and PAP2) and anthocyanin synthesis genes under CAS condition. Stars represent significantly changed (FDR < 0.05) genes compared to parental line. (F) Venn diagram and Gene Ontology (GO) enrichment of differentially expressed genes (DEGs) in mutant rosettes from CAS and potting conditions to identify signaling responses.

### The Schengen pathway conveys soil nitrogen status to shoots independent of Casparian strip status

Under CAS conditions, both WTs and *myb36-2* plants showed signs of stress-induced anthocyanin accumulation in the leaves, whereas MYB36Loop rosettes remained green (Fig. 5A). The *sgn3-3* mutant phenocopied the MYB36Loop line in terms of repressed anthocyanin accumulation (Fig. 5A), which supports that this stress response is SGN-dependent as these two mutants have opposing CS status. On CAS, all rosettes contained only minute amounts of nitrate (Fig. 5D) while the level of other nutrients such as phosphate and sulfate showed only minor changes in the mutants under both soil conditions (Fig. S5B). As nitrogen starvation can induce anthocyanin accumulation [26, 27], we tested if the lack of this response in the MYB36Loop and *sgn3-3* plants under CAS conditions was due to impaired N-signaling. For this, we performed a transcriptional analysis of rosettes from *myb36-2*, MYB36Loop #5 as well as the *sgn3-3* mutant on different soil types. Indeed, specifically under CAS conditions, MYB36Loop and *sgn3-3* rosettes had induced expression of nitrogen-signaling genes belonging to the NITRATE-INDUCIBLE, GARP-TYPE (NIGT) family [21, 22] as well as the LATERAL BOUNDARY DOMAIN 37, 38 and 39 (LBD37,38 and 39) [28] (Fig. 5E). Under nitrogen-sufficient conditions, LBD37-39 repress expression of the anthocyanin master regulators PRODUCTION OF ANTHOCYANIN PIGMENT 1 and 2 (PAP1 and PAP2) [28-30]. These responses were also observed in MYB36Loop and *sgn3-3*, along with repression of PAP target genes in the anthocyanin biosynthetic pathway (Fig. 5E) [31]. Combined, this strongly indicates that under CAS conditions, an active SGN pathway is required to convey N-starvation responses from roots to shoots. In line with this, among the 253 DEGs specifically repressed in MYB36Loop and *sgn3-3* under CAS conditions (Fig. S5C), we found GO terms related to anthocyanin and flavonoid production as well as response to N-starvation (Fig. 5F). Moreover, within the same subset of genes we found that processes related to leaf senescence, salt stress and salicylic acid were specifically enriched under CAS conditions (Fig. 5F). As these were induced in *myb36-2*, this further supports that specifically SGN responses were repressed in the MYB36Loop shoots. However, and quite remarkably, under potting soil conditions, there were very few genes present in the overlap between the DEGs repressed in MYB36Loop and *sgn3-3*, but not differentially expressed in *myb36-2* rosettes (Fig. 5F). As this indicates that signals sent from roots via the SGN pathway are soil-status dependent, we further investigated the N-responses observed on CAS. Upon watering with a nutrient solution containing Ca(NO_3_)_2_, the parental lines and *myb36-2* plants showed a strong growth promotion, which was accompanied by a decrease in anthocyanin accumulation when compared to mock treatment (Fig. 6A-C). However, and in strong support of the defective N-signaling, the growth-promoting effect was not occurring in the *sgn3-3* or MYB36Loop rosettes (Fig. 6A-C). Combined, we conclude that under soil conditions that are close to natural, the SGN pathway is essential to convey nitrogen status of the soil to the shoots. Thus, rather than functioning as a CS survaillance mechanism, this signaling pathway is likely to function as an overall integrator of CS and soil status into long-distance signaling to the shoots (Fig. 7).

**Figure 6.**
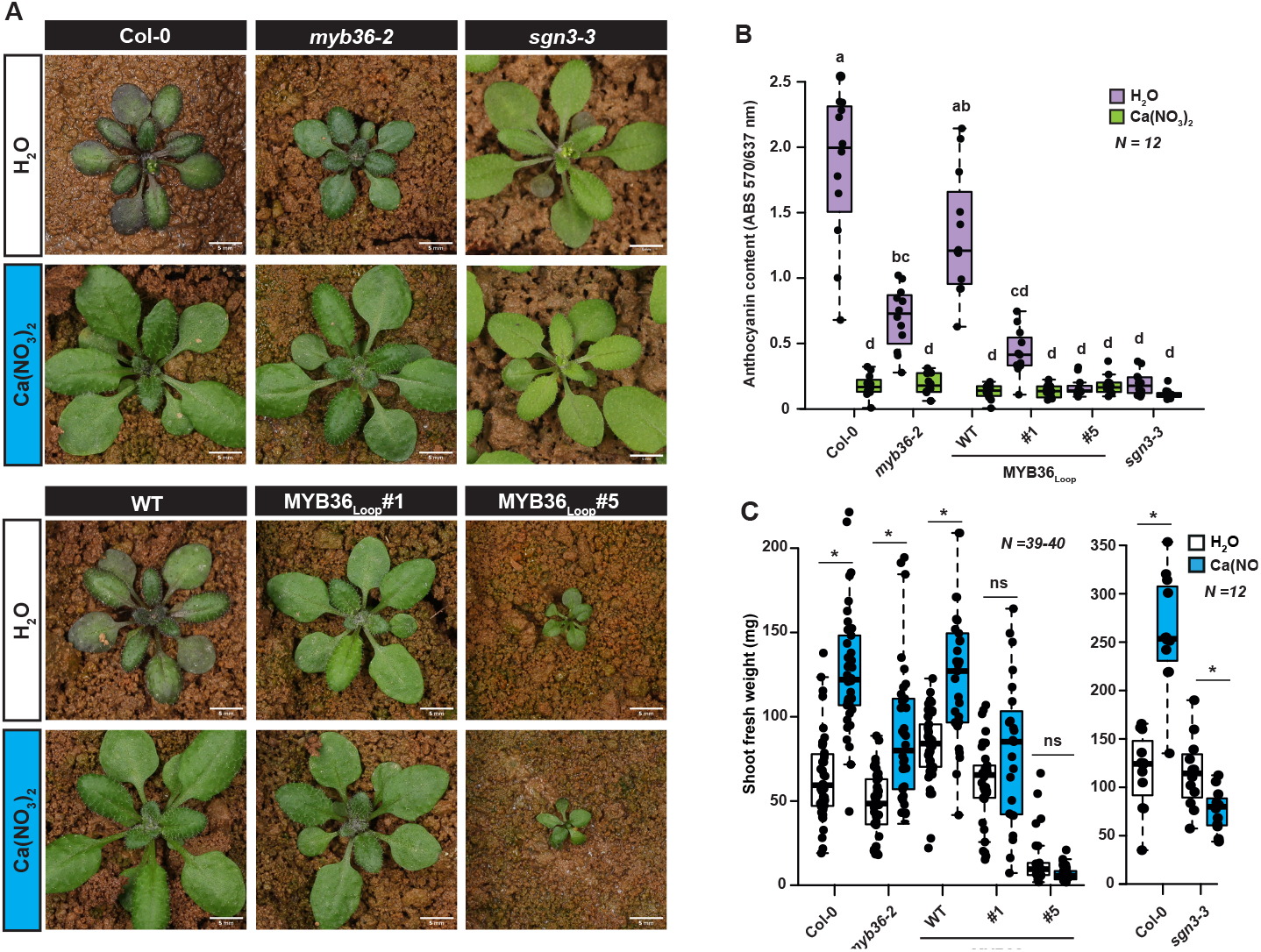
Growth analysis of mutants grown on agricultural soil under a nitrogen fertilization scheme. (A) Four-week-old rosettes from different genotypes grown for one week on ½ MS conditions and transferred to Cologne agricultural soil (CAS) for three weeks and watered with water (H2O) or a CalciniteTM solution containing nitrate (Ca(NO3)2). Scale bars represent 5 mm. (B) Anthocyanin content of 4-week-old rosettes from CAS watered with mock (H2O) or nitrate (Ca(NO3)2). (C) Shoot fresh weight of rosettes from CAS watered with H2O or nitrate (Ca(NO3)2). Asterisks depict significant changes between H2O and nitrate treatments according to a two-sided Student’s t-test (P < 0.05). ns; not significant. For box-plots, center line in box indicates median, dots represent data, limits represent upper and lower quartiles, and whiskers maximum and minimum values. Letters depict statistical difference of one-way ANOVA with Holm-Sidak-adjusted posthoc t-test (P < 0.05).

**Figure 7.**
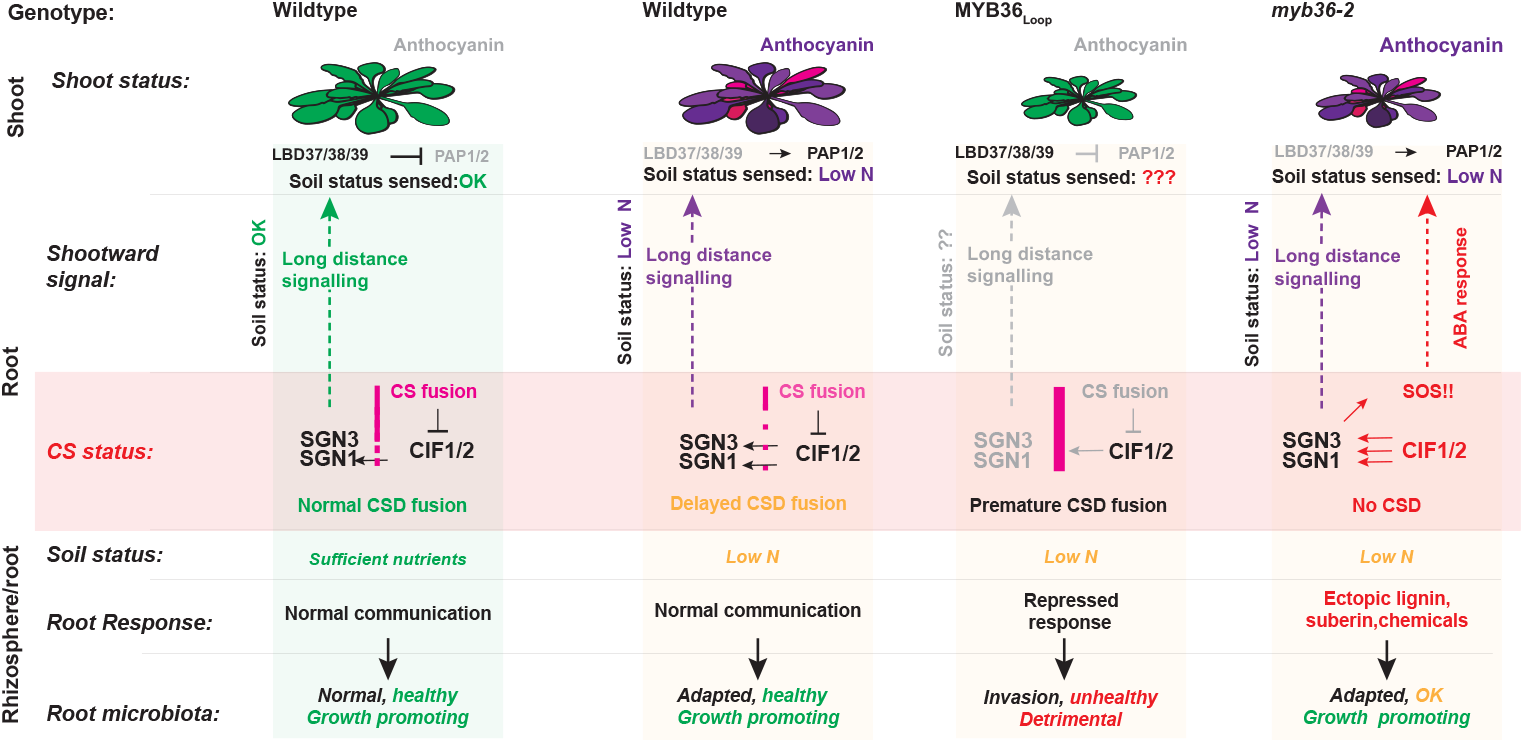
Model for Casparian strip-related signaling across different soil conditions. In roots Casparian strip (CS) fusion depends on activation of the Schengen pathway (SGN1 and SGN3) via diffusion of the SGN3 ligands known as Casparian strip integrity factor (CIF) peptides from the stele across the endodermis. In our model, this feature additionally provides information to the shoot and allows correct communication with the rhizosphere. Upon low nitrogen status in the soil, the CS establishment is delayed and gives rise to an increased diffusion of CIF and thereby a quantitative difference in SGN signaling to the shoot. Upon increased CS formation in the MYB36Loop lines, this signaling pathway is overridden and no long-distance signaling can occur, which blocks activation of the SGN pathway and makes the shoot unable to sense the soil environment. Moreover, increased CS formation disturbs the ability of the root to communicate with the rhizosphere. In case of disturbed CS formation (i.e. myb36 KO) the lack of CS induces an ABA-dependent SOS response in roots which is propagated independently to the shoot and capable of compensating for the rhizosphere communication. Thus, it is likely that a separate signaling mechanism downstream of SGN3 capable of facilitating long-distance signaling exists.

## Discussion

Positive feedback loops have been identified in genetic networks that control secondary cell wall formation [32] and play a role in reinforcing robust differential outputs in epidermal development [33]. These are tightly regulated to avoid run-away expression. Thus, an artificial positive transcriptional feedback loop puts strain on the endogenous regulatory network. This can manifest either in form of transcriptional co-factor depletion or compensatory repressive mechanisms such as silencing, that eventually must weaken the exponential expression encouraged by the feedback. If this is not the case, the plant will exhaust all resources in order to facilitate continuous loop function. With this in mind, despite our success in creating a feedback loop that exaggerates CS formation in the endodermis, it is likely that unspecific side effects occur and caution should be taken when using such constructs. Despite this, we find that this loop manifested a number of systemic consequences in plants, which can be directly ascribed to the increase in barrier formation and/or the decoupling of the SGN system.

Combined, these point towards a model where under complex soil conditions, the interplay between root barriers and the associated signaling serves as a mechanism to integrate soil status into their shoot nutrient homeostasis. This provides an updated long-distance signaling model where the Schengen pathway is essential for integration of the CS status into systemic signaling that informs aboveground plant parts of the nutritional environment (Fig.7). In this model, spatial changes in the onset of CS establishment — which can occur as a response to external stimuli such as low nitrogen conditions — result in changed CIF diffusion from the stele. This is then read out as a quantitative change in SGN activation which allows fine-tuned integration of CS status into downstream signaling mechanisms (Fig 7). However, upon disturbed CS formation (i.e. *myb36-2*), this system appears to be still active, suggesting that the route for signaling in case of hyperactivated SGN is different than the nutrient responsive system. Schengen system bears resemblance to a number of receptor kinase systems related to development and immunity, where downstream signaling is conveyed by different phosphorylation pathways via cytoplasmic kinases [34]. It is therefore possible that the downstream signaling from SGN3 is facilitated by different downstream kinases, of which only SGN1/PBL15 has so far been identified.

While a number of studies have given comprehensive insights into the impact of these barriers on mineral accumulation in plant tissues, very little is known of barrier-associated changes in nitrogen accumulation and responses. We here provide direct and quantitative evidence that beneficial nitrogen-related growth responses require correct coordination of CS-SGN signaling in roots. As a microbial disbalance was evident in MYB36Loop expressing plants, it is plausible that the observed lack of nitrogen-induced growth promotion was due to an inability to establish a healthy root microbiome. This is supported by recent evidence in rice (Oryza sativa) where the bacterial communities play an important role in nitrogen uptake and usage [35]. Long-distance nitrogen-signaling has also been extensively studied in legumes due to their ability to form nitrogen-fixing relationships with rhizobacteria [36]. In these systems, upwards movement of peptides convey signals from roots to shoots, causing responses under low nitrogen availability [37]. While these systems are less characterized in Arabidopsis, it is plausible that components of the long-distance nitrogen signaling mechanisms may be downstream targets of the SGN system. Thus, under nitrogen-scarce conditions, activation of the SGN system in the root provides a mechanism that allows the plant to integrate root and rhizosphere status in shoot responses. Moreover, in context of the disturbed root microbial communities, the lack of nitrogen-induced growth promotion in the MYB36Loop lines implies that this is a consequence of an inhibited ability to exploit microbe-mediated benefits.

In summary, our work provides a direct mechanistic link between CS establishment, shoot physiology and microbial community establishment in the rhizosphere. Our work demonstrates that in an agriculturally-relevant root environment, timing of the CS onset is essential for establishing a physiological homeostasis in local as well as in distal tissues. This provides an updated insight into how plants coordinate communication between their above- and belowground parts and emphasizes a so-far overlooked phenomenon where root barriers are important for integrating soil information into overall physiological responses.

## Acknowledgments

All authors thank Bart Boesten, and Ila Rouhara for technical s upport. Sabine Ambrosius and the Biocenter MS Platform Cologne for measurements of mineral composition. Ton Timmers and the Central Microscopy facility (CeMic) are thanked for microscopy aid. Aristeidis Stamatakis and his greenhouse team at MPIPZ are thanked for help with plant growth. We moreover thank Meike Burow, Niko Geldner, Magdalena Marek, Sebastian Samwald, Lioba Ruger and Marc Somssich for insightful comments on the manuscript. Joop Vermeer and his lab are thanked for comments to our preprint version of this work. We also thank Satoshi Fujita and Hiroko Uchida for graphical support. TGA would further like to thank BDR and APM for their support in form of liquid aid at timely intervals. Funding: Research in the lab of TGA is supported by the Sofja Kovalevskaja programme from the Alexander von Humboldt foundation and the Max Planck Society. Research in the lab of SK is supported by the Deutsche Forschungsgemeinschaft (DFG) under Germany’s Excellence Strategy – EXC 2048/1 – project 390686111. KW is funded by DFG Priority Programme SPP2125 DECRyPT.

## Author contributions

Conceptualization: TGA. Methodology: DS, KW, NK, UN, SK, PZ, TGA. Investigation: DS, SR, KW, NK, UN, SK, TGA. Visualization: DS, KW and TGA. Funding acquisition: TGA, SK. Main writing: TGA. All authors read and commented on the final version of the manuscript.

### Competing interests

All authors declare that they have no competing interests.

### Data and materials availability

RNA-seq raw reads and amplicon sequencing raw reads generated in this study have been deposited at National Center for Biotechnology Information under BioProject ID PRJNA940103 and PRJNA961040.

## Materials and Methods

### Plant growth

Arabidopsis thaliana ecotype Columbia-0 transgenic and mutant lines were used to perform experiments. Seeds were kept for 2 days at 4°C in the dark for stratification. For nutrient starvation experiments plants were grown under 16 h light at 21°C and 8 h dark at 19°C vertically on ½ MS medium without sucrose but with distinct nutrient compositions. In all media, the pH was buffered to 5.8 using 10 mM 2-(N-morpholino) ethanesulfonic acid (MES). The following premixed media were used for nutrient starvation assays: Caisson MSP11, MSP44 or MSP21. Low nitrogen conditions were obtained by Caisson MSP21 + 0.037 mM KNO3 + 0.037 mM NH4NO3. For fresh weight measurements, shoots and roots of five plants were harvested after 14 days (3 days on ½ MS +11 days on the specified condition). For Cologne agriculture soil (CAS) [25] and potting soil (Blumavis Mini Tray MIM800) experiments, 4-5 plants were grown in 9×9 cm square pots in a climate-controlled greenhouse, at 21°C under a 16 h light/8 h dark regime. For CAS experiments pots were placed on top of a capillary mattress in a tray, and tap water or calcinitTM solution (0.651 g/L, calcinitTM composition: 14.4% nitrate, 1% ammonium, 26% calcium oxide) (Yara, Germany) was added. For potting soil experiments, soil was moisturized by submerging the pot bottom in water regularly. For gnotobiotic experiments with SynCom inoculation, the FlowPot system was used as previously described [24]. The SynCom strains are part of the At-RSPHERE collection [38] and are listed in Table S5.

### Cloning

To generate the endodermis-specific feedback loop expression constructs, the coding sequence of MYB36 [8] was Gateway-cloned into a pDONR221 entry vector using BP clonase II (Invitrogen) according to manufacturer’s description. Together with previously generated P4L1r pDONR entry vectors containing the pCASP1 sequence [3], this was recombined using LR-clonase II (Invitrogen) into a FastRed selection-containing destination vector (pED97) [39]. The final construct was transformed into pCASP1::CASP1-GFP and other backgrounds using the floral dip method and selected using FastRed selection.

### Staining procedures

All staining procedures were done using ClearSee staining [40, 41]. Briefly, plants were fixed in 3 mL 1 x PBS containing 4% p-formaldehyde for 1 hour at room temperature and washed twice with 3 mL 1 x PBS. Following fixation, the seedlings were cleared in 3 mL ClearSee solution (10% xylitol, 15% sodium deoxycholate and 25% urea in water) under gentle shaking. After overnight clearing, the solution was exchanged to new ClearSee solution containing 0.2% Basic Fuchsin and 0.1% Calcofluor White for lignin and cell wall staining respectively. The dye solution was removed after overnight staining and rinsed once with fresh ClearSee solution. The samples were washed for 30 min with gentle shaking followed by overnight incubation in ClearSee solution before imaging. Suberin staining of Arabidopsis roots was performed as previously described [41]. In case of combined suberin and lignin staining the procedure was according to [42]. Briefly, vertically grown 5-day old seedlings were incubated solution of Fluorol Yellow 088 (Sigma) (0.01%, in lactic acid) and incubated for 30 min at 70°C. The stained seedlings were rinsed shortly in water and transferred to a freshly prepared solution of Aniline blue (0.5%, in water) for counterstaining. Propidium iodide (PI) assays were done as described [43]. PI staining seedlings were washed for 2-3 min in water and transferred to a chambered cover glass (Thermo Scientific), and imaged either using Confocal laser scanning (CLSM) microscopy or epifluorescence microscopy.

### Microscopy

All confocal images were taken using a Zeiss LSM 980 system. Fluorophore settings were ex 488nm, em 505-550 nm for GFP and Fluorol Yellow 088, ex 594 nm, em 610-650 nm for mCherry, ex 561 nm, em 580-600 nm for Basic fuchsin, ex 405 nm em 420-430 nm for Calcofluor White. Airyscan images were taken using the 4Y multiplex setting. FY staining and PI analysis were done on a Zeiss Axiozoom V16 system (GFP filtercube ex: 470 nm/40 em:525/50 bs: 500). For PI TX2 filtercube ex: 560 nm/40 em:645/75 bs: 595 nm).

### Transmission electron microscopy

For quantification of the Casparian strip width by TEM, 6-day-old pCASP1::CASP1-GFP and MYB36Loop#5 seedlings were placed in small Petri dishes filled with 0.05 M MOPS buffer supplemented with 0.1% Tween20 and 10 mM cerium chloride and incubated with gentle agitation at room temperature. After 30 min, the CeCl3-containing MOPS buffer was replaced by 2.5% glutaraldehyde in 0.05 M phosphate buffer (v/v), pH 7.2, and seedlings were gently agitated for one hour. Subsequently, seedlings were transferred into glass vials filled with 1% osmium tetroxide (EMS, #19150) in 0.05 M phosphate buffer (w/v), pH 7.2, supplemented with 1.5% potassium ferrocyanide, for another hour. After three rinses in water, seedlings were embedded into 2% low melting temperature agarose and three fragments were sampled from the distal 1.5 cm-long part of the root. Root fragments were then dehydrated with a series of ethanol, gradually transferred into acetone and embedded into Araldite 502/Embed 812 resin (EMS, #13940) using the EMS Poly III embedding machine (EMS, #4444). Ultrathin sections (≈70 nm) were cut at specific distances from the root tip with a Reichert-Jung Ultracut E and collected on Formvar-coated copper slot grids [44]. After staining with 0.1% potassium permanganate in 0.1 N H2SO4 for one minute (w/v) followed by 0.5% uranyl acetate in water (w/v) for 10 minutes and lead citrate for 15 min, sections were examined with a Hitachi H-7650 TEM operating at 100 kV and equipped with an AMT XR41-M digital camera. For immunogold detection of CASP1-GFP in MYB36Loop roots were high-pressure frozen in a Leica EM HPM 100 high-pressure freezer between two large aluminum specimen carriers (ø 4.6 mm) enclosing a cavity of 150 µm depth filled with ½ MS medium. After freeze substitution in the Leica EM AFS2 freeze substitution device using 0.5% uranyl acetate in acetone (w/v), bringing samples from -85°C to -20°C over seven days, samples were transferred to ethanol and gradually embedded into LR White resin (Plano GmbH, R1281) at -20°C over six days with constant agitation. Samples were polymerized in pure LR White resin with UV light for 24 h at -20°C and 24 h at 0°C. Ultramicrotomy was performed as described above with the exception that sections were collected on Formvar-coated gold slot grids. Immunogold labelling of CASP1-GFP was carried out according to [45] using a 1:5 dilution of rat monoclonal anti-GFP 3H9 (Chromotek) and a 1:20 dilution of goat anti-rat IgG conjugated to 10 nm colloidal gold particles (British Biocell International). Sections were stained with potassium permanganate and uranyl acetate (no lead citrate) and imaged as described above.

### Transcriptomics

For transcriptomic analysis on roots, Col-0, *myb36-2*, pCASP1::CASP1-GFP (WT), MYB36Loop#1 and MYB36Loop#5 seedlings were grown on standard ½ MS agar medium for seven days. Whole roots were harvested and immediately frozen in liquid nitrogen. RNA was extracted using a TRIzol (Invitrogen)-adapted ReliaPrep RNA extraction kit (Promega) [39]. For transcriptomic analysis on CAS and potting soil grown shoots (set 1), Col-0, *myb36-2*, WT, and MYB36Loop#5 seedlings were grown on standard solid ½ MS medium for 7 days, then transferred to CAS soil or potting soil. Note that transcriptomic analysis on CAS and potting soil grown *sgn3-3* shoots was an independent experiment (set 2), WT and *sgn3-3* (expressing pCASP1::CASP1-GFP) samples were prepared as mentioned above. Whole rosette leaves of 4-week-old plants were harvested and immediately frozen in liquid nitrogen. RNA extractions were performed as mentioned above. RNA quality was determined using a Bioanalyzer 2100 system (Agilent Technologies, USA). Library preparation and paired-end 150 bp sequencing were conducted by Novogene (Cambridge, UK). Approximately 40 million raw reads were generated per sample. RNA-seq raw reads of *sgn3-3* roots were acquired from [11, 14], raw reads of CIF2 treated Col-0 roots dataset (48 hours after CIF2 treatment) were acquired from [11]. All raw reads were preprocessed using fastp (v0.22.0) [46]. Filtered high-quality reads were mapped to A. thaliana TAIR10 reference genome with Araport 11 annotation (Phytozome genome ID: 447) using HISAT2 (v2.2.1), and counted using featureCounts from the Subread package (v2.0.1) [46]. All statistical analyses were performed using R (v4.1.2) (https://www.R-project.org/). Read counts were transformed to cpm (counts per million) using edgeR package [47]. Lowly expressed (less than 0.5 x total sample number 19 for ½ MS roots, 26 for CAS shoots set 1, 8 for CAS shoots set 2, 16 for potting shoots set 1, 8 for potting shoots set 2, 6 for [14] 9 for [11]) cpm over all samples, and at least 1 cpm in n samples (6 for ½ MS roots, 8 for CAS shoots set 1, 2 for CAS shoots set 2, 5 for potting shoots set 1, 2 for potting shoots set 2, 2 for [14], 3 for [11]) genes were removed from the analysis. Differentially expressed genes (DEGs) were identified by pairwise comparisons using the glmFit function in edgeR package [47] with absolute fold change more than 2 and a false discovery rate (FDR) corrected P-value less than 0.05. Note that for root transcriptome analysis, MYB36Loop#1 and MYB36Loop#5 samples were compiled as one genotype, and compared with WT. Gene Ontology (GO) term enrichment analysis was performed using Metascape [48]. Only GO terms with adjusted P value (q-value) lower than 0.05 were considered as significantly enriched. Heatmaps were generated using pheatmap package (https://cran.r-project.org/web/packages/pheatmap/index.html) or ggplot2 package [49]. Upset plots were generated using UpSetR package.

### ICP-MS and IC analysis

The total mineral content was determined by inductively-coupled plasma mass spectrometry (ICP-MS) following the method described in [50]. Approximately 5 mg of homogenized dried plant material were digested using 500 μL of 67% (w/w) HNO3 overnight at room temperature and subsequently placed in a 95°C water bath for 30 min or until the liquid was completely clear. After cooling to room temperature, the samples were placed on ice and 4.5 mL of deionized water was carefully added to the tubes. The samples were centrifuged at 4°C at 2,000 g for 30 min and the supernatants were transferred to new tubes. The elemental concentration was determined using Agilent 7700 ICP-MS (Agilent Technologies) [50]. Inorganic anion (nitrate, phosphate, and sulfate) levels were measured by ion chromatography, as described in [51]. Approximately 10 mg of dried plant material was homogenized in 1 mL deionized water, shaken for 1 h at 4°C, and subsequently heated at 95°C for 15 min. The anions were determined by the Dionex ICS-1100 chromatography system and separated on a Dionex IonPac AS22 RFIC 4× 250 mm analytic column (Thermo Scientific, Darmstadt, Germany), using 4.5 mM Na2CO3/1.4 mM NaHCO3 as running buffer [51]. To compare the shoot ionomes between different genotypes under different growth conditions, a principal coordinate analysis (PCoA) was performed with a Bray– Curtis dissimilarity index calculated using vegdist() function in the R vegan package (https://github.com/jarioksa/vegan). To assess the variations explained by growth conditions (CAS vs. potting soil), a PERMANOVA testing was performed, based on the Bray–Curtis dissimilarity index, using adonis2() function from the R vegan package with 999 permutations. To assess the statistical difference of shoot ionomes between different genotypes within each growth condition, a pairwise comparison (PERMANOVA) was performed, based on the Bray–Curtis dissimilarity index, using the pairwise. adonis() function in the R pairwise Adonis package (https://github.com/pmartinezarbizu/pairwiseAdonis) with 999 permutations, followed by Benjamini-Hochberg post hoc testing.

### Community profiling

For community profiling on CAS grown root samples, plants were grown on standard ½ MS agar medium for 7 days, then transferred to CAS soil and grown for three weeks. Three to four plants from the same pot were collected as one replicate. The whole root systems of three to four plants were carefully isolated, rhizosphere and root compartments were harvested as previously described [24]. All samples (including unplanted soil samples) were transferred to Lysing Matrix E tubes (FastDNA Spin Kit for Soil, MP Biomedicals), frozen in liquid nitrogen and stored at −80 °C for further processing. DNA was isolated using a FastDNA Spin Kit for Soil (MP Biomedicals), then subjected to bacterial and fungal community profiling, and absolute quantification of bacterial and fungal load, as previously described [52]. For community profiling on roots derived from FlowPot experiments, roots of all plants in one FlowPot were harvested as one sample, cleaned from attaching soil particles, and processed further like described above. Library preparation for amplicon sequencing was performed as described previously [53]. Briefly, multiplexing of samples was performed by double-indexing with barcoded forward and reverse oligonucleotides for 16S rRNA gene or fungal ITS amplification. Samples from the FlowPot experiment were sequenced on the Illumina Novaseq platform at Novogene, UK. Samples of the greenhouse experiment in CAS soil were sequenced on our in-house Illumina MiSeq platform. Amplicon data processing was performed as previously described [24]. Shortly, the reads were demultiplexed by QIIME2 plugin ‘cutadapt demux-paired’ [54]. Demultiplexed reads were merged with Flash2 [54]. For synthetic communities, the merged reads were fed into Rbec (https://github.com/PengfanZhang/Rbec) to profile the microbial communities. For natural communities, reads were fed into DADA2 to generate the ASV table. Chimeras were detected using Vsearch [55] and excluded from the downstream analysis. Taxonomic classification of ASVs were conducted by the Bayesian classifier trained on the SILVA (Ver. 138) and Unite (Ver. 8) for bacteria and fungi respectively [56, 57]. All the ASVs assigned to mitochondria and chloroplasts were eliminated from the ASV table.

### Anthocyanin content measurement

Measurement of anthocyanin content in rosettes grown under CAS or standard potting soil conditions was performed as previously described [58]. Absorbances at 530 and 637 nm were measured using Tecan Infinite 200 PRO plate reader.

---

**Figure S1.**
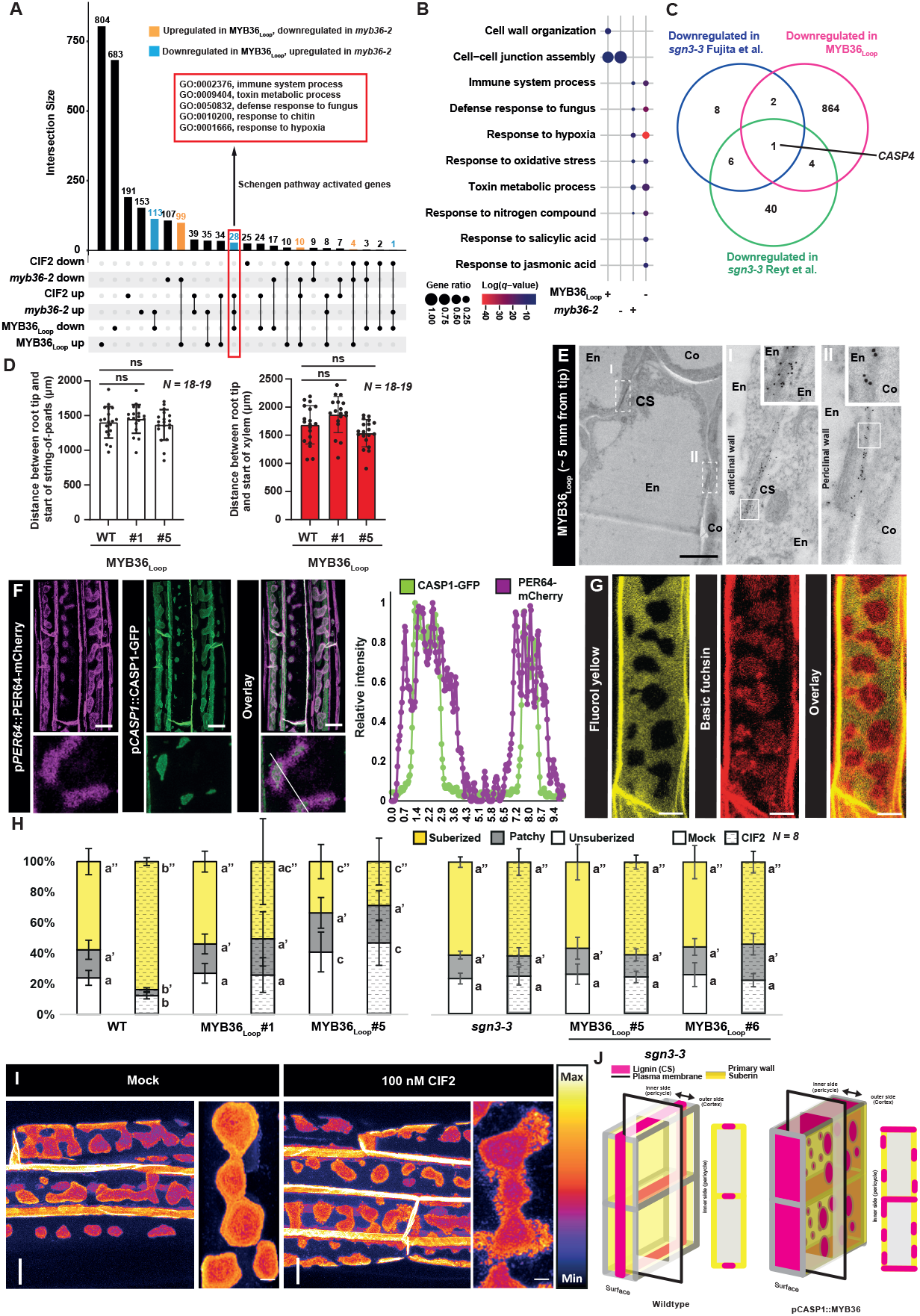
(A) Upset plot showing the number of differentially expressed genes (DEGs) in MYB36_Loop_ and *myb36-2* roots grown on standard ½ MS agar medium, comparing with the DEGs in Col-0 roots treated with CIF2 from [11]. Orange bars represent genes upregulated in MYB36_Loop_, downregulated in myb36-2; blue bars represent genes downregulated in MYB36_Loop_, upregulated in *myb36-2*. Red rectangle highlights Schengen pathway activated genes, with GO terms enriched. (B) Gene Ontology (GO) term enrichment of DEGs. Note that GO terms enriched in MYB36_Loop_ downregulated genes are largely overlapping with the GO terms enriched in Schengen pathway activated genes in (A). Gene ratio represents the number of DEGs of GO term divided by the total genes in the GO term. + represents upregulated genes, - represents downregulated genes. (C) Venn diagram showing the overlap of downregulated genes in MYB36_Loop_r oots, downregulated genes in *sgn3-3* roots from [14] and downregulated genes in Col-0 roots treated with CIF2 [11]. (D) Bar plots showing the distance between root tip and start of string-of-pearls CSD region (left graph) and the distance between root tip and start of xylem (right graph). Plants were grown on standard ½ MS agar medium for 8 days. Data are mean ± SD. Statistical significance of differences with the parental line (WT) was determined using a two-tailed Student’s t test. ns; not significant. (E)Transmission electron microscopy (TEM) micrograph of 7-day-old anti-GFP immunogold-labeled MYB36_Loop_ #5 roots. Scale bar represents 500 nm. Co; Cortex, CS; Casparian strip, En: Endodermis. (F) Maximum projection of a confocal image stack of 7-day-old MYB36_Loop_ plants expressing p*PER64*::PER64-mCherry and p*CASP1*::CASP1-GFP. Line in overlay depicts the transect used for relative intensity measurements. (G) Endodermal cells of plants expressing MYB36_Loop_ stained with Basic Fuchsin and Fluorol yellow. (H) Measurement of suberization pattern under mock or 100 nM CIF2 conditions. (I) Top-view maximum projection of 7-days old MYB36_Loop_#5 plants treated with 100 nM CIF2 for 24h before fixing in Clearsee and imaged using a confocal setup. Scale bars represent 10 µm. For boxplots, the center line in the box indicates the median, dots represent data, the box limits represent the upper and lower quartiles, and the whiskers represent the maximum and minimum values. Different letters depict statistical difference in a one-way ANOVA analysis with Tukey’s test (P < 0.05).

**Figure S2.**
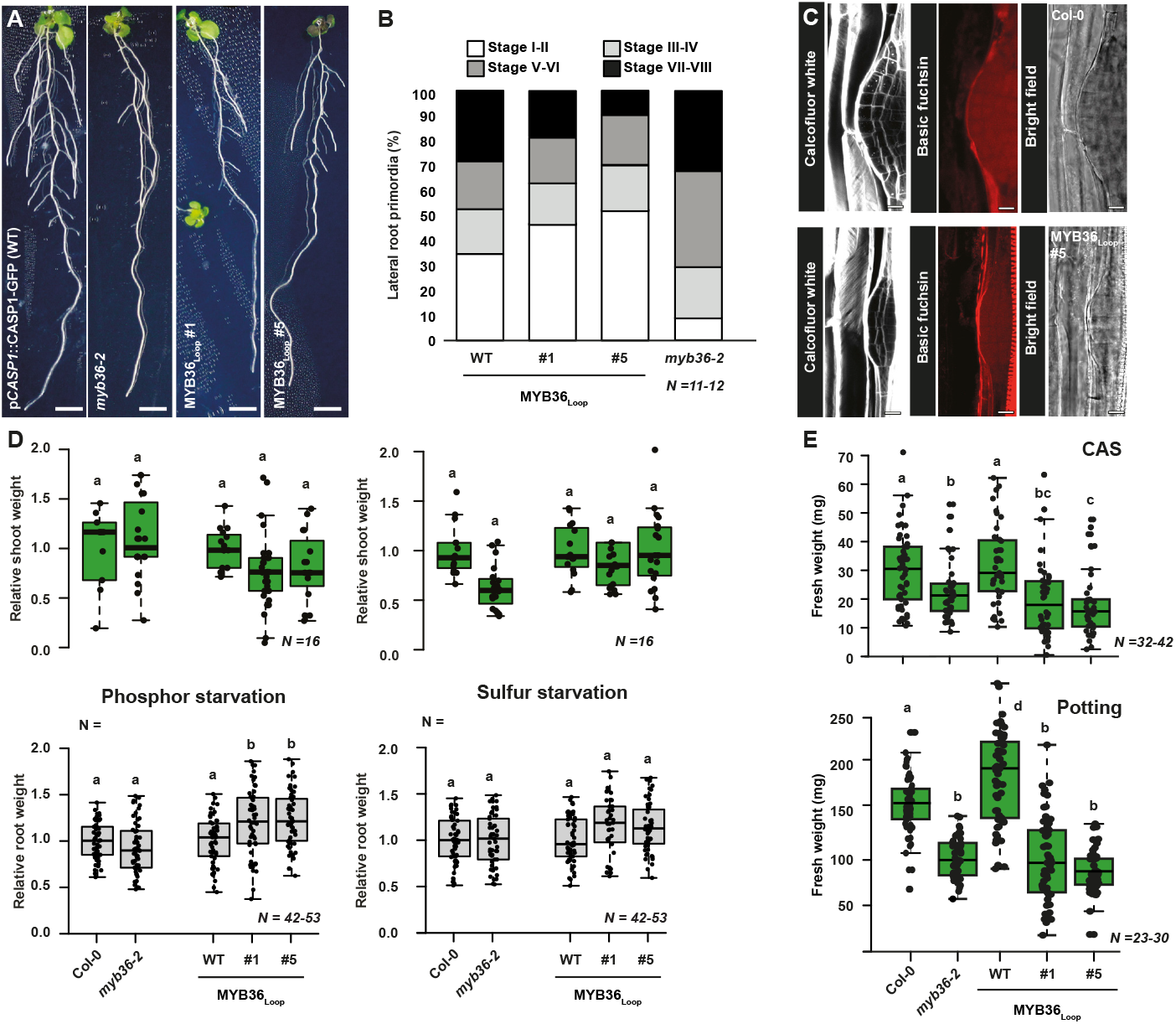
(A) 14-day-old seedlings grown on standard ½ MS agar medium. Scale bars represent 5 mm. (B) The distribution of different stages of Lateral root primordia (LRP) of 8-day-old plants grown on standard ½ MS agar medium. (C) LRP of 8-day-old Col-0 (upper graph) and MYB36_Loop_#5 (lower graph) roots grown on standard ½ MS agar medium, stained with cell wall dye Calcofluor White and the lignin-specific dye Basic Fuchsin. Scale bars represent 10 µm. (D) Measurement of shoot (upper graph) and root (lower graph) fresh weight of 2-week-old plants grown under phosphor (left) or sulfur (right) starvation conditions. The weight was normalized to the changes in the corresponding parental background (Col-0 for *myb36-2* and p*CASP1*::CASP1-GFP for MYB36_Loop_ lines). (E) Measurement of shoot fresh weight of four-week-old plants grown under CAS (upper graph) or standard potting soil (lower graph) conditions. The growth was normalized to the changes in the corresponding parental background (Col-0 for *myb36-2* and p*CASP1*::CASP1-GFP (WT) for MYB36_Loop_ lines). For boxplots, the center line in the box indicates the median, dots represent data, the box limits represent the upper and lower quartiles, and the whiskers represent the maximum and minimum values. Different letters depict statistical difference in a one-way ANOVA analysis with Tukey’s test (P < 0.05).

**Figure S3.**
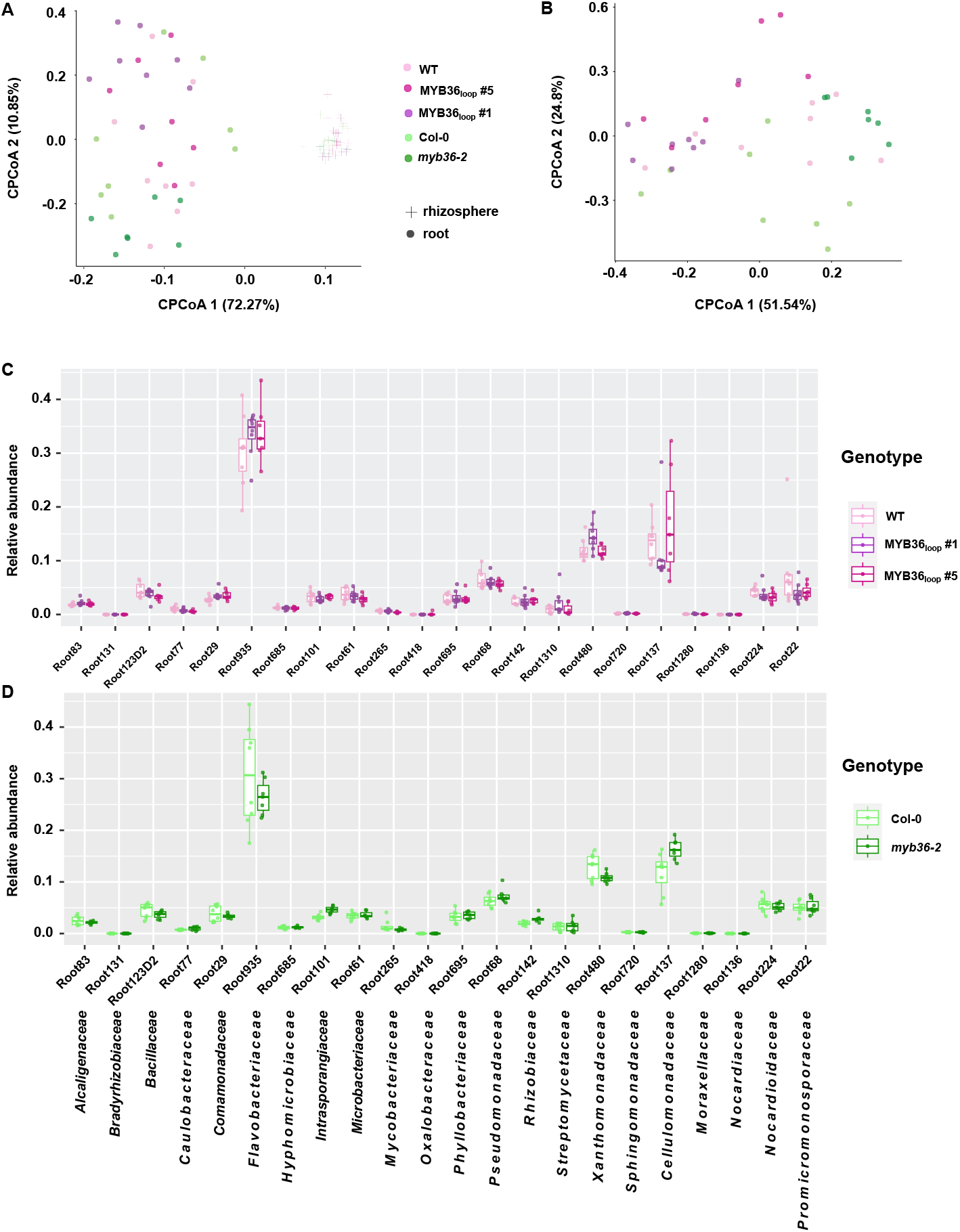
(A) Constrained Principal Coordinates Analysis (CPCoA) of Bray-Curtis dissimilarities of 16S amplicon reads from root and rhizosphere samples (41.7% of variance explained by genotype x compartment, P=0.001, PERMANOVA). (B) CPCoA of root samples only (17.8% of variance explained by genotype, P=0.003, PERMANOVA). (C and D) Relative abundance of individual SynCom strains (IDs shown as x-axis labels, and corresponding bacterial families below) in root samples of p*CASP1*::-CASP1-GFP (WT), MYB36_Loop_#1 and MYB36_Loop_#5 (C), and of Col-0 and *myb36-2* (D).

**Figure S4.**
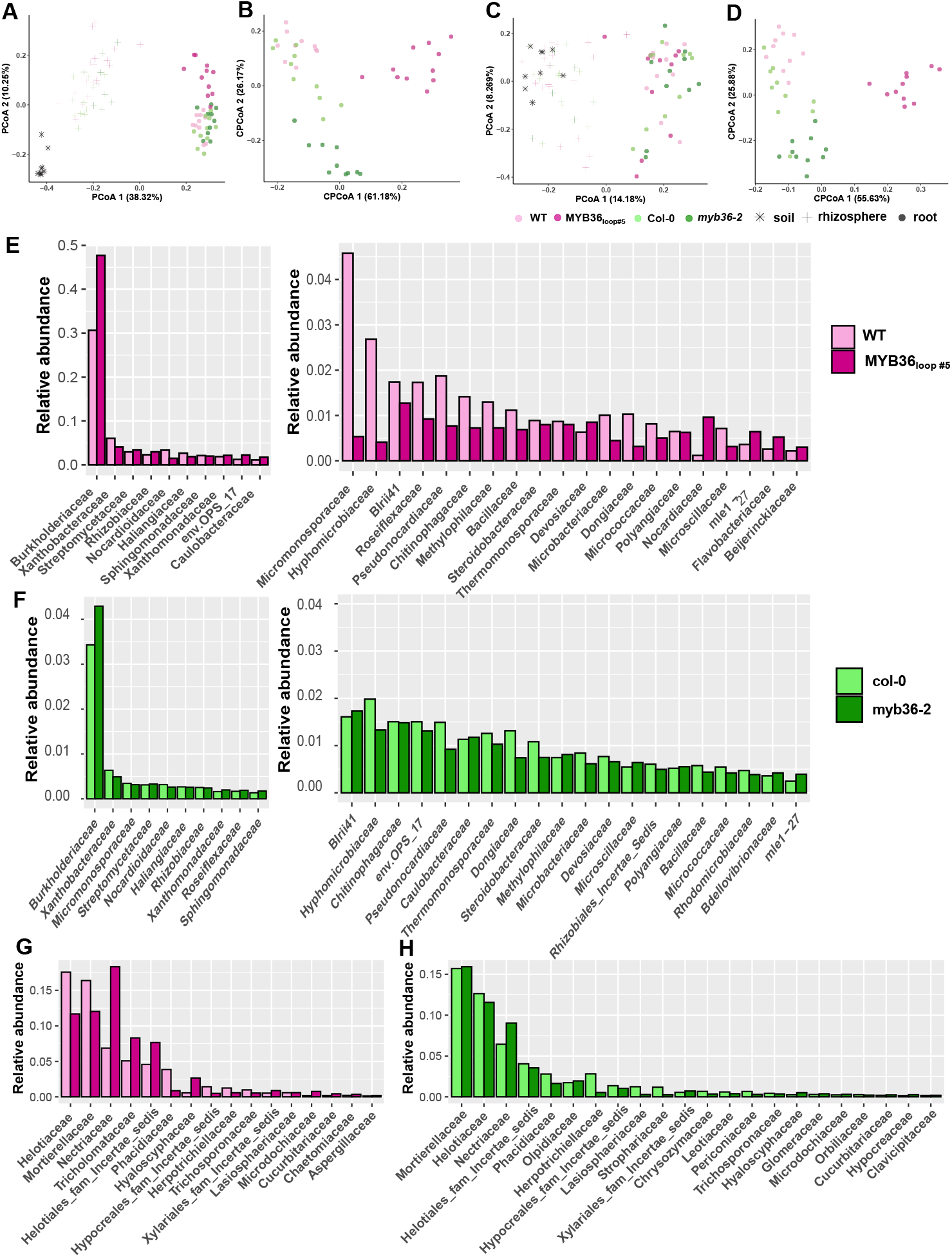
(A and C) PCoA of Bray-Curtis dissimilarities of bacterial 16S amplicon reads (A) and fungal ITS amplicon reads (C) from p*CASP1*::CASP1-GFP (WT), MYB36_Loop_ #5, Col-0, and *myb36-2* root and rhizosphere samples as well as unplanted soil samples. (B and D) Constrained PCoA of the same root samples, where 15.9% and 9.3% of the variance are explained by genotype for bacteria (B) and fungi (D), respectively (P=0.001, PERMANOVA). (E-H) Rank plot of median relative abundances of abundant bacterial (E and F) and fungal (G and H) families in MYB36_Loop_ #5 and *myb36-2* roots and their corresponding parental lines.

**Figure S5.**
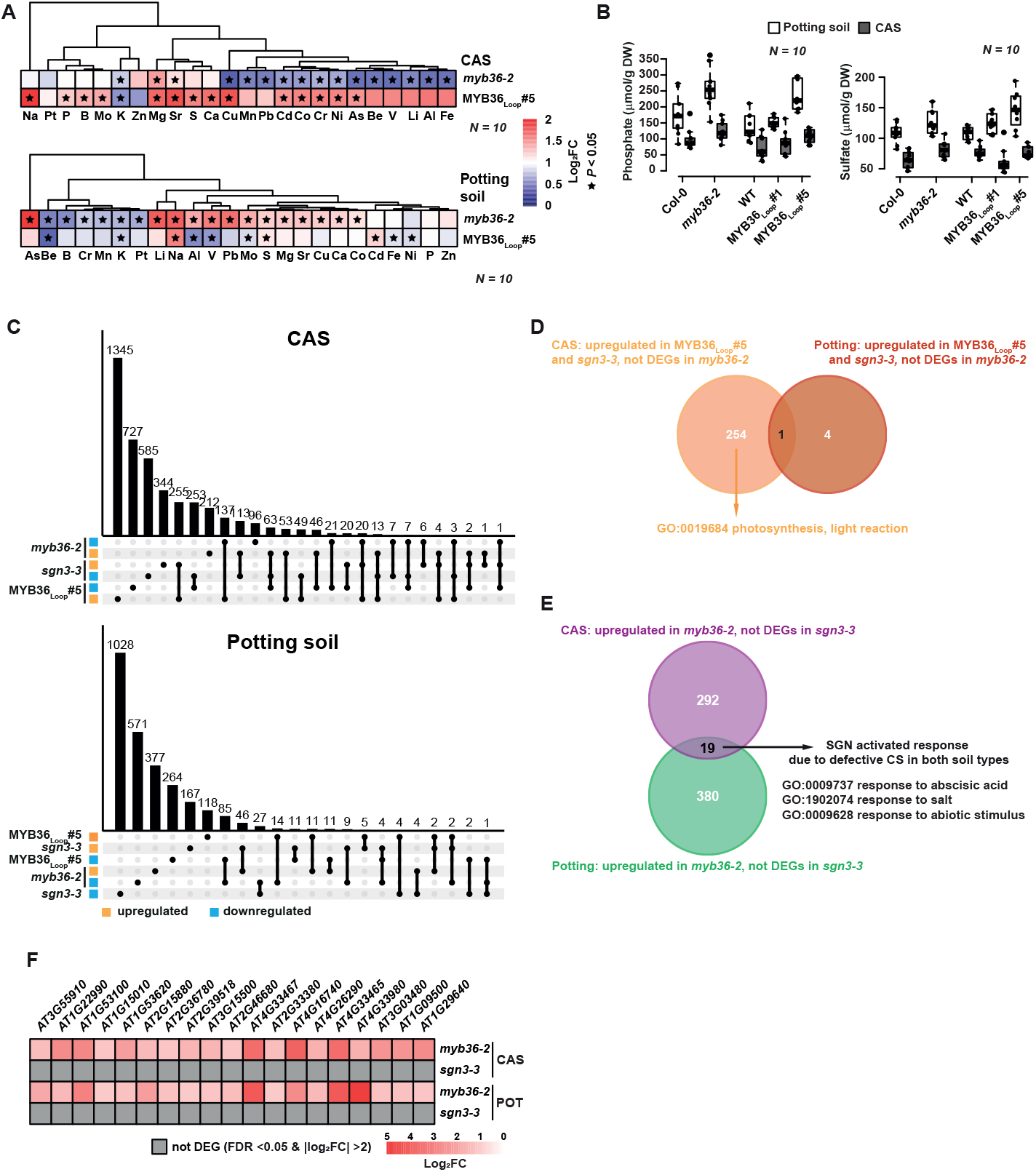
Heatmap of 24 element contents in 4-week-old MYB36_Loop_#5 and *myb36-2* rosettes grown under CAS (upper) or standard potting soil (lower) conditions after normalization to the corresponding parental line. Star symbols represent significant differences with respective wild-type line determined using a two-tailed Student’s t test (P < 0.05). (B) Measurement of phosphate (left) and sulfate (right) content in 4-week-old rosettes grown under CAS or standard potting soil conditions. (C) Upset plots showing the number of DEGs in MYB36_Loop_ #5, *sgn3-3* and *myb36-2* rosettes grown under CAS (upper) or standard potting soil (lower) conditions. Orange: upregulated; blue: downregulated. (D) Venn diagram depicting the overlap of DEGs in rosettes grown under CAS and standard potting conditions and the GO terms specifically enriched among the 254 upregulated DEGs in MYB36_Loop_#5 and *sgn3-3* rosettes, but not DEGs in *myb36-2* rosettes under CAS condition. (E) Venn diagram depicting the overlap of upregulated DEGs in *myb36-2*, but not DEGs in *sgn3-3* rosettes between CAS and standard potting condition, and the GO terms enriched. The overlap represents 19 genes activated by the Schengen (SGN) pathway due to defective Casparian strip (CS) formation in both soil conditions. (F) Heatmap indicating the 19 genes activated by SGN pathway due to defective Casparian strip in both soil types. Grey boxes represent genes that do not meet the DEG threshold. For boxplots, the center line in the box indicates the median, dots represent data, the box limits represent the upper and lower quartiles, and the whiskers represent the maximum and minimum values.

## Notes

### Competing Interest Statement

The authors have declared no competing interest.

### Summary of Updates

We included transcriptional data on the sgn3-3 mutant as well as phenotypes when grown on CAS. We included new data on behavior of MYB36Loop in the esb1 mutant. The text has been edited and updated to clarify findings We also included figure 7, which contains a descriptive model that puts our findings in context

## References

1. Geldner, N. (2013). The Endodermis. Annual Review of Plant Biology 64, 531–558.

2. Barberon, M., and Geldner, N. (2014). Radial Transport of Nutrients: The Plant Root as a Polarized Epithelium. Plant Physiology 166, 528–537.

3. Roppolo, D., De Rybel, B., Dénervaud Tendon, V., Pfister, A., Alassimone, J., Vermeer, J.E., Yamazaki, M., Stierhof, Y.D., Beeckman, T., and Geldner, N. (2011). A novel protein family mediates Casparian strip formation in the endodermis. Nature 473, 380–383.

4. Hosmani, P.S., Kamiya, T., Danku, J., Naseer, S., Geldner, N., Guerinot, M.L., and Salt, D.E. (2013). Dirigent domain-containing protein is part of the machinery required for formation of the lignin-based Casparian strip in the root. Proc Natl Acad Sci U S A 110, 14498–14503.

5. Baxter, I., Hosmani, P.S., Rus, A., Lahner, B., Borevitz, J.O., Muthukumar, B., Mickelbart, M.V., Schreiber, L., Franke, R.B., and Salt, D.E. (2009). Root Suberin Forms an Extracellular Barrier That Affects Water Relations and Mineral Nutrition in Arabidopsis. PLOS Genetics 5, e1000492.

6. Lee, Y., Rubio Maria C., Alassimone, J., and Geldner, N. (2013). A Mechanism for Localized Lignin Deposition in the Endodermis. Cell 153, 402–412.

7. Liberman, L.M., Sparks, E.E., Moreno-Risueno, M.A., Petricka, J.J., and Benfey, P.N. (2015). MYB36 regulates the transition from proliferation to differentiation in the <i>Arabidopsis</i> root. Proceedings of the National Academy of Sciences 112, 12099–12104.

8. Kamiya, T., Borghi, M., Wang, P., Danku, J.M.C., Kalmbach, L., Hosmani, P.S., Naseer, S., Fujiwara, T., Geldner, N., and Salt, D.E. (2015). The MYB36 transcription factor orchestrates Casparian strip formation. Proceedings of the National Academy of Sciences 112, 10533–10538.

9. Pfister, A., Barberon, M., Alassimone, J., Kalmbach, L., Lee, Y., Vermeer, J.E.M., Yamazaki, M., Li, G., Maurel, C., Takano, J., et al. (2014). A receptorlike kinase mutant with absent endodermal diffusion barrier displays selective nutrient homeostasis defects. eLife 3, e03115.

10. Doblas, V.G., Smakowska-Luzan, E., Fujita, S., Alassimone, J., Barberon, M., Madalinski, M., Belkhadir, Y., and Geldner, N. (2017). Root diffusion barrier control by a vasculature-derived peptide binding to the SGN3 receptor. Science 355, 280–284.

11. Fujita, S., De Bellis, D., Edel, K.H., Köster, P., Andersen, T.G., Schmid-Siegert, E., Dénervaud Tendon, V., Pfister, A., Marhavý, P., Ursache, R., et al. (2020). SCHENGEN receptor module drives localized ROS production and lignification in plant roots. The EMBO Journal 39, e103894.

12. Fujita, S. (2021). CASPARIAN STRIP INTEGRITY FACTOR (CIF) family peptides – regulator of plant extracellular barriers. Peptides 143, 170599.

13. Salas-González, I., Reyt, G., Flis, P., Custódio, V., Gopaulchan, D., Bakhoum, N., Dew, T.P., Suresh, K., Franke, R.B., Dangl, J.L., et al. (2021). Coordination between microbiota and root endodermis supports plant mineral nutrient homeostasis. Science 371.

14. Reyt, G., Ramakrishna, P., Salas-González, I., Fujita, S., Love, A., Tiemessen, D., Lapierre, C., Morreel, K., Calvo-Polanco, M., Flis, P., et al. (2021). Two chemically distinct root lignin barriers control solute and water balance. Nature Communications 12, 2320.

15. Wang, F.-L., Tan, Y.-L., Wallrad, L., Du, X.-Q., Eickelkamp, A., Wang, Z.-F., He, G.-F., Rehms, F., Li, Z., Han, J.-P., et al. (2021). A potassium-sensing niche in Arabidopsis roots orchestrates signaling and adaptation responses to maintain nutrient homeostasis. Developmental Cell 56, 781–794.e786.

16. Wang, P., Calvo-Polanco, M., Reyt, G., Barberon, M., Champeyroux, C., Santoni, V., Maurel, C., Franke, R.B., Ljung, K., Novak, O., et al. (2019). Surveillance of cell wall diffusion barrier integrity modulates water and solute transport in plants. Scientific Reports 9, 4227.

17. Andersen, T.G., Molina, D., Kilian, J., Franke, R.B., Ragni, L., and Geldner, N. (2021). Tissue-autonomous phenylpropanoid production is essential for establishment of root barriers. Current Biology 31, 965–977. e

18. Schalk, M., Cabello-Hurtado, F., Pierrel, M.A., Atanossova, R., Saindrenan, P., and Werck-Reichhart, D. (1998). Piperonylic acid, a selective, mechanism-based inactivator of the trans-cinnamate 4-hydroxylase: A new tool to control the flux of metabolites in the phenylpropanoid pathway. Plant Physiol 118, 209–218.

19. Fernández-Marcos, M., Desvoyes, B., Manzano, C., Liberman, L.M., Benfey, P.N., del Pozo, J.C., and Gutierrez, C. (2017). Control of Arabidopsis lateral root primordium boundaries by MYB36. New Phytologist 213, 105–112.

20. Konishi, M., and Yanagisawa, S. (2013). Arabidopsis NIN-like transcription factors have a central role in nitrate signalling. Nature communications 4, 1617.

21. Kiba, T., Inaba, J., Kudo, T., Ueda, N., Konishi, M., Mitsuda, N., Takiguchi, Y., Kondou, Y., Yoshizumi, T., and Ohme-Takagi, M. (2018). Repression of nitrogen starvation responses by members of the Arabidopsis GARP-type transcription factor NIGT1/HRS1 subfamily. The Plant Cell 30, 925–945.

22. Araus, V., Vidal, E.A., Puelma, T., Alamos, S., Mieulet, D., Guiderdoni, E., and Gutiérrez, R.A. (2016). Members of BTB gene family of scaffold proteins suppress nitrate uptake and nitrogen use efficiency. Plant Physiology 171, 1523–1532.

23. Guo, Y., Wang, Y., Chen, H., Du, Q., Wang, Z., Gong, X., Sun, Q., and Li, W.X. (2023). Nitrogen supply affects ion homeostasis by modifying root Casparian strip formation through the miR528-LAC3 module in maize. Plant Commun, 100553.

24. Wippel, K., Tao, K., Niu, Y., Zgadzaj, R., Kiel, N., Guan, R., Dahms, E., Zhang, P., Jensen, D.B., Logemann, E., et al. (2021). Host preference and invasiveness of commensal bacteria in the Lotus and Arabidopsis root microbiota. Nature Microbiology 6, 1150–1162.

25. Harbort, C.J., Hashimoto, M., Inoue, H., Niu, Y., Guan, R., Rombolà, A.D., Kopriva, S., Voges, M., Sattely, E.S., Garrido-Oter, R., et al. (2020). Root-Secreted Coumarins and the Microbiota Interact to Improve Iron Nutrition in Arabidopsis. Cell Host Microbe 28, 825–837. e826.

26. Chalker-Scott, L. (1999). Environmental significance of anthocyanins in plant stress responses Photochemistry and Photobiology 70.

27. Diaz, C., Saliba-Colombani, V., Loudet, O., Belluomo, P., Moreau, L., Daniel-Vedele, F., Morot-Gaudry, J.-F., and Masclaux-Daubresse, C. (2006). Leaf yellowing and anthocyanin accumulation are two genetically independent strategies in response to nitrogen limitation in Arabidopsis thaliana. Plant and Cell Physiology 47, 74–83.

28. Rubin, G., Tohge, T., Matsuda, F., Saito, K., and Scheible, W.-R.d. (2009). Members of the LBD Family of Transcription Factors Repress Anthocyanin Synthesis and Affect Additional Nitrogen Responses in Arabidopsis The Plant Cell 21, 3567–3584.

29. Rowan, D.D., Cao, M., Lin-Wang, K., Cooney, J.M., Jensen, D.J., Austin, P.T., Hunt, M.B., Norling, C., Hellens, R.P., and Schaffer, R.J. (2009). Environmental regulation of leaf colour in red 35S: PAP1 Arabidopsis thaliana. New Phytologist 182, 102–115.

30. Li, N., Wu, H., Ding, Q., Li, H., Li, Z., Ding, J., and Li, Y. (2018). The heterologous expression of Arabidopsis PAP2 induces anthocyanin accumulation and inhibits plant growth in tomato. Functional & integrative genomics 18, 341–353.

31. Dooner, H.K., Robbins, T.P., and Jorgensen, R.A. (1991). Genetic and developmental control of anthocyanin biosynthesis. Annual review of genetics 25, 173–199.

32. Taylor-Teeples, M., Lin, L., de Lucas, M., Turco, G., Toal, T.W., Gaudinier, A., Young, N.F., Trabucco, G.M., Veling, M.T., Lamothe, R., et al. (2015). An Arabidopsis gene regulatory network for secondary cell wall synthesis. Nature 517, 571–575.

33. Kang, Y.H., Kirik, V., Hulskamp, M., Nam, K.H., Hagely, K., Lee, M.M., and Schiefelbein, J. (2009). The MYB23 gene provides a positive feedback loop for cell fate specification in the Arabidopsis root epidermis. Plant Cell 21, 1080–1094.

34. Zhang, J., Li, W., Xiang, T., Liu, Z., Laluk, K., Ding, X., Zou, Y., Gao, M., Zhang, X., Chen, S., et al. (2010). Receptor-like Cytoplasmic Kinases Integrate Signaling from Multiple Plant Immune Receptors and Are Targeted by a Pseudomonas syringae Effector. Cell Host & Microbe 7, 290–301.

35. Zhang, J., Liu, Y.-X., Zhang, N., Hu, B., Jin, T., Xu, H., Qin, Y., Yan, P., Zhang, X., Guo, X., et al. (2019). NRT1.1B is associated with root microbiota composition and nitrogen use in field-grown rice. Nature Biotechnology 37, 676–684.

36. Gautrat, P., Laffont, C., Frugier, F., and Ruffel, S. (2021). Nitrogen Systemic Signaling: From Symbiotic Nodulation to Root Acquisition. Trends in Plant Science 26, 392–406.

37. Roy, S., and Müller, L.M. (2022). A rulebook for peptide control of legume–microbe endosymbioses. Trends in Plant Science 27, 870–889.

38. Bai, Y., Müller, D.B., Srinivas, G., Garrido-Oter, R., Potthoff, E., Rott, M., Dombrowski, N., Münch, P.C., Spaepen, S., Remus-Emsermann, M., et al. (2015). Functional overlap of the Arabidopsis leaf and root microbiota. Nature 528, 364–369.

39. Andersen, T.G., Naseer, S., Ursache, R., Wybouw, B., Smet, W., De Rybel, B., Vermeer, J.E.M., and Geldner, N. (2018). Diffusible repression of cytokinin signalling produces endodermal symmetry and passage cells. Nature 555, 529–533.

40. Kurihara, D., Mizuta, Y., Sato, Y., and Higashiyama, T. (2015). ClearSee: a rapid optical clearing reagent for whole-plant fluorescence imaging. Development 142, 4168–4179.

41. Ursache, R., Andersen, T.G., Marhavý, P., and Geldner, N. (2018). A protocol for combining fluorescent proteins with histological stains for diverse cell wall components. Plant J 93, 399–412.

42. Sexauer, M., Shen, D., Schön, M., Andersen, T.G., and Markmann, K. (2021). Visualizing polymeric components that define distinct root barriers across plant lineages. Development 148.

43. Naseer, S., Lee, Y., Lapierre, C., Franke, R., Nawrath, C., and Geldner, N. (2012). Casparian strip diffusion barrier in <i>Arabidopsis</i> is made of a lignin polymer without suberin. Proceedings of the National Academy of Sciences 109, 10101–10106.

44. Moran, D.T., and Rowley, J.C. (1987). 1 – Biological Specimen Preparation for Correlative Light and Electron Microscopy.

45. Viñegra de la Torre, N., Vayssières, A., Obeng-Hinneh, E., Neumann, U., Zhou, Y., Lázaro, A., Roggen, A., Sun, H., Stolze, S.C., Nakagami, H., et al. (2022). FLOWERING REPRESSOR AAA+ ATPase 1 is a novel regulator of perennial flowering in Arabis alpina. New Phytologist 236, 729–744.

46. Chen, S., Zhou, Y., Chen, Y., and Gu, J. (2018). fastp: an ultra-fast all-in-one FASTQ preprocessor. Bioinformatics 34, i884–i890.

47. Robinson, M.D., McCarthy, D.J., and Smyth, G.K. (2009). edgeR: a Bioconductor package for differential expression analysis of digital gene expression data. Bioinformatics 26, 139–140.

48. Zhou, Y., Zhou, B., Pache, L., Chang, M., Khodabakhshi, A.H., Tanaseichuk, O., Benner, C., and Chanda, S.K. (2019). Metascape provides a biologist-oriented resource for the analysis of systems-level datasets. Nature Communications 10, 1523.

49. Wickham, H. (16 June 2016). ggplot2:Elegant Graphics for Data Analysis, Volume XVI, (Springer Cham).

50. Almario, J., Jeena, G., Wunder, J., Langen, G., Zuccaro, A., Coupland, G., and Bucher, M. (2017). Root-associated fungal microbiota of nonmycorrhizal <i>Arabis alpina</i> and its contribution to plant phosphorus nutrition. Proceedings of the National Academy of Sciences 114, E9403–E9412.

51. Dietzen, C., Koprivova, A., Whitcomb, S.J., Langen, G., Jobe, T.O., Hoefgen, R., and Kopriva, S. (2020). The Transcription Factor EIL1 Participates in the Regulation of Sulfur-Deficiency Response. Plant Physiol 184, 2120–2136.

52. Hou, S., Thiergart, T., Vannier, N., Mesny, F., Ziegler, J., Pickel, B., and Hacquard, S. (2021). A microbiota–root–shoot circuit favours Arabidopsis growth over defence under suboptimal light. Nature Plants 7, 1078–1092.

53. Durán, P., Thiergart, T., Garrido-Oter, R., Agler, M., Kemen, E., Schulze-Lefert, P., and Hacquard, S. (2018). Microbial Interkingdom Interactions in Roots Promote Arabidopsis Survival. Cell 175, 973–983.e914.

54. Bolyen, E., Rideout, J.R., Dillon, M.R., Bokulich, N.A., Abnet, C.C., Al-Ghalith, G.A., Alexander, H., Alm, E.J., Arumugam, M., Asnicar, F., et al. (2019). Reproducible, interactive, scalable and extensible microbiome data science using QIIME 2. Nature Biotechnology 37, 852–857.

55. Rognes, T., Flouri, T., Nichols, B., Quince, C., and Mahé, F. (2016). VSEARCH: a versatile open source tool for metagenomics. PeerJ 4, e2584.

56. Quast, C., Pruesse, E., Yilmaz, P., Gerken, J., Schweer, T., Yarza, P., Peplies, J., and Glöckner, F.O. (2013). The SILVA ribosomal RNA gene database project: improved data processing and web-based tools. Nucleic Acids Res 41, D590–596.

57. Nilsson, R.H., Larsson, K.-H., Taylor, A.F S., Bengtsson-Palme, J., Jeppesen, T.S., Schigel, D., Kennedy, P., Picard, K., Glöckner, F.O., Tedersoo, L., et al. (2018). The UNITE database for molecular identification of fungi: handling dark taxa and parallel taxonomic classifications. Nucleic Acids Research 47, D259–D264.

58. Nakata, M., and Ohme-Takagi, M. (2014). Quantification of Anthocyanin Content. Bio-protocol 4, e1098.

